# Assessing the capacity of high-resolution commercial satellite imagery for grapevine downy mildew detection and surveillance in New York state

**DOI:** 10.1101/2023.11.10.566469

**Authors:** Kathleen Kanaley, David B. Combs, Angela Paul, Yu Jiang, Terry Bates, Kaitlin M. Gold

## Abstract

Grapevine downy mildew (GDM), caused by the oomycete *Plasmopara viticola,* can cause 100% yield loss and vine death under conducive conditions. Growers currently rely on frequent fungicide applications for control, but this practice has led to widespread resistance. Rapid remote detection and surveillance of GDM outbreaks would enable precision pesticide applications to target effective but resistance-prone fungicides where and when most needed, while relying on less resistance-prone protectants elsewhere. High resolution commercial satellite platforms offer the opportunity to track rapidly spreading diseases like GDM over large, heterogeneous fields. Here, we investigate the capacity of PlanetScope (3 m) and SkySat (50 cm) imagery for season-long GDM detection and surveillance. A team of trained scouts rated GDM severity and incidence in two acres of Chardonnay grapevines in Geneva, NY, USA in June-August of 2020, 2021, and 2022. Satellite imagery acquired within 72 hours of scouting was processed to extract single-band reflectance and vegetation indices (VIs). Random forest models trained on spectral bands and VIs derived from both image datasets could classify areas of high and low GDM incidence and severity with maximum accuracies of 0.88 (SkySat) and 0.94 (PlanetScope). However, we do not observe significant differences between VIs of high and low damage classes until late July-early August. We identify cloud cover, image co-registration, and low spectral resolution as key challenges to operationalizing satellite-based GDM surveillance. This work establishes the capacity of spaceborne multispectral sensors to detect late-stage GDM and outlines steps towards incorporating satellite remote sensing in grapevine disease surveillance systems.

## I. Introduction

Effective plant disease control is crucial to ensure food and nutritional security for growing populations in a changing climate. Disease prevention and management is most effective at the earliest stages of pathogen establishment before outbreaks become severe. However, the scale at which modern agriculture is conducted, over hundreds of thousands of acres, makes it logistically infeasible for growers to monitor all acres of production for early signs of disease. The primary barrier to early disease detection is the labor and time required to scout for infections, which has inspired great interest in the use of remote sensing for both disease detection and surveillance. Disease detection differs from disease surveillance in that “detection” refers to disease identification at a particular location and point in time, while “surveillance” means consistent, ongoing monitoring of plant disease status. Disease detection is a critical first step to building operational disease surveillance systems with remote sensing. Disease detection technologies aim to accurately identify a specific cause of crop stress, while surveillance systems seek to provide this via repeated measures of the same location(s) over time to direct resources to areas that deviate from a healthy baseline.

The need for disease surveillance is particularly important for grapevine downy mildew (GDM), an endemic foliar necrotizing disease caused by the obligate hemi-biotroph oomycete *Plasmopara viticola*. GDM can result in complete crop loss within a single season if left unmanaged (Wilcox, Gubler, and Uyemoto 2015). GDM control is achieved primarily through foliar fungicide applications, which can offer significant protection if fungicides are applied before infections occur. However, fungicides have limited curative action once *P. viticola* sufficiently colonizes grapevine tissue (Gessler, Pertot, and Perazzolli 2011). *Plasmopara viticola*’s rapid polycyclic infection cycle enables it to propagate quickly throughout the vineyard, making fungicide resistance development a great concern. Multiple rounds of sexual reproduction can occur within a single growing season as spores from distinct individuals infect the same plant host. If a vineyard’s pathogen population is exposed to repeated applications of fungicides with the same mode of action (MOA), it becomes more likely that individuals resistant to that MOA will dominate the gene pool as their susceptible counterparts are eliminated (Campbell et al. 2021; Toffolatti et al. 2018; Gisi and Sierotzki 2008; Corio-Costet 2012).

GDM poses a great challenge to grape growers in the northeastern United States in particular, who have a limited number of specialty “oomicides” registered to manage the disease. Resistance to some of the major FRAC groups, including carboxylic acid amide (CAA) and quinone outside inhibitor (QOI) fungicides has already been documented in the Finger Lakes AVA (Sharma et al. 2022), and surveys of *P. viticola* populations in other cool climate grape-growing regions suggest that resistance to these products will become more widespread if populations continue to be exposed to these chemistries (Santos et al. 2020; Feng and Baudoin 2018; Campbell et al. 2021; Corio-Costet 2012; Sharma et al. 2022). An effective and scalable method for early GDM detection and operationalized surveillance would enable grape growers to better strategize where, when, and most importantly, *what* chemical control measures to apply. If growers had an accurate way to monitor GDM establishment and progression, they could rely on less effective but less resistance-prone chemistries for preventative control, switching to higher efficacy but more resistance-prone chemistries when warning systems are triggered. The ability to rapidly assess GDM-induced losses is important for evaluating the efficacy of control measures that have already been applied. Coupled with fungicide application records, remote disease surveillance could aid in evaluating the prevalence of resistant pathogen populations at a regional to national scale. Disease surveillance can also contribute to estimating yield losses and generating geographically explicit year-to-year comparisons of disease impact on yields and/or fruit quality.

For decades, satellite remote sensing has been used for plant disease detection and surveillance across a range of agricultural and natural ecosystems (Junming Wang et al. 2010; Jing Wang et al. 2020; Z. Wang et al. 2016; Khanal et al. 2020; Nagarajan et al. 1984; Nutter et al. 2002). Until recently, the utility of satellite imagery for agricultural disease detection and surveillance has been hampered by lack of adequate spatial, temporal, and spectral resolution to monitor the target vegetation type (Nilsson 1995; Jackson 1986; Moran, Inoue, and Barnes 1997). Prior to the 2010s, lengthy revisit times and the coarse spatial resolution of most satellite platforms made accurate surveillance impractical, especially for diseases like GDM which can spread exponentially in a matter of days. Freely available data from space agencies, such as Landsat 9 and Sentinel-2 imagery, is too spatially coarse to detect disease symptoms in individual grapevine panels. Further challenges arise in vineyards and trellised cropping systems where rows of plants are separated by inter-rows of soil, cover crops, and other vegetation. This within-field heterogeneity makes it difficult to attribute changes in reflectance to the target crop.

Recent engineering innovations and advances in satellite constellation design have improved the temporal and radiometric consistency as well as scalability of spaceborne sensing platforms. These platforms include DigitalGlobe’s World View fleet (0.5-1.2 m spatial resolution, 8-38 spectral bands), the Pleiades constellation operated by AirBus Defense (2 m, 4 bands), and Planet Lab’s SuperDove (3 m, 8 bands) and SkySat-C (0.5 m, 4 bands) fleets (Yang 2018; Frazier and Hemingway 2021; Saunier et al. 2022). The SuperDoves operate on a sun-synchronous orbit with a 24-hr revisit time and offer global coverage. Imagery is available as a harmonized product that is radiometrically consistent with coincident Sentinel-2 acquisitions (Tu et al. 2022; Kim et al. 2022). Planet’s SkySat C fleet offers imagery at 50 cm spatial resolution in four multispectral bands (red, green, blue, NIR) and is currently available as a tasked product with revisit time dependent on the area of interest and contract specifications (Saunier et al. 2022; “Planet Imagery Product Specifications” 2022).

The proliferation of the commercial high resolution Earth observation industry has reinvigorated research interest in spaceborne ecological surveillance. SkySat and PlanetScope imagery has been used for a range of ecosystem and agricultural applications, including biomass estimation, vegetation classification, plant disease detection, quantifying evapotranspiration, and soil moisture mapping (Kharel et al. 2023; Guo et al. 2022; Szabó et al. 2021; Shi et al. 2018; Baloloy et al. 2018; Du et al. 2022; Raza et al. 2020; Aragon et al. 2021). The frequent revisit times and fine spatial scale of these platforms enables monitoring diverse specialty crops, including grapevine, despite their smaller, spatially heterogeneous fields (Helman et al. 2018; Meyers et al. 2020). However, most prior work on grapevine remote sensing has aimed to detect water stress, with few studies addressing other abiotic parameters such as nitrogen content, yield, and fruit composition (Giovos et al. 2021). Reports on proximal and remote sensing of grapevine diseases are dominated by near-surface platforms and increasingly by hyperspectral cameras (both proximal and airborne) (Naidu et al. 2009; Oerke, Herzog, and Toepfer 2016; MacDonald et al. 2016; Bendel et al. 2020; Gao et al. 2020; Lacotte et al. 2022; Sawyer et al. 2023; di Gennaro et al. 2016; Matese et al. 2022; Cséfalvay et al. 2009; Galvan et al. 2023). Most studies aim to detect disease at a single point in time, while season-long, operational surveillance systems remain largely unexplored.

To our knowledge, there are no published efforts to use high resolution satellite imagery to detect and track grapevine downy mildew. Despite the lack of investigation, GDM is a suitable target for detection and surveillance via satellite-derived multispectral imagery. Foliar signs and symptoms include sporulation on the leaf underside and chlorosis on the adaxial leaf surface, which progresses to abaxial leaf necrosis when left unchecked (Koledenkova et al. 2022). Of these symptoms, chlorosis and reduction of green leaf area are the most relevant indicators for remote disease detection. Photosynthesis is downregulated in infected leaf tissue, with a consequent decrease in chlorophyll pigments (Koledenkova et al. 2022). Chlorophyll absorbs solar radiation strongly in the red (∼620-670 nm) and blue (∼420-470 nm) regions of the electromagnetic spectrum and reflects more light in the green (∼520-570 nm). By reducing chlorophyll content, GDM causes increased reflectance in visible wavelengths no longer absorbed for photosynthesis. Additionally, while the parenchyma and mesophyll scatter near-infrared light (∼870-1200 nm) in healthy plants, in vines undergoing *P. viticola* colonization the pathogen breaks down the cellular structures that normally contribute to NIR reflectance (Knipling 1970; Koledenkova et al. 2022). We hypothesize that spectral divergence of healthy vs. infected plant tissue driven by these physiological changes can enable GDM detection and surveillance via multispectral satellite imagery.

Here, we investigate the application of Planet Lab’s PlanetScope and SkySat imagery for GDM detection and surveillance in a 2-acre Chardonnay vineyard in Geneva, New York, USA. We set out to: 1) determine whether PlanetScope and SkySat imagery can be used to detect GDM in a vineyard; 2) evaluate performance of satellite disease detection and surveillance across multiple growing seasons at the same site to assess the robustness of remotely sensed disease indicators across time; 3) test supervised machine learning for predicting GDM damage using satellite-derived spectral features; and 4) identify potential limitations and future opportunities for using commercial Earth observations in GDM surveillance systems.

## II. Materials and Methods

### Study location

The study was conducted in a 30-year-old Cornell pathology research vineyard in Geneva, New York, USA (42°52’41.8’’ N, 77°0’54.8’’). The study block contains 20 rows of *Vitis vinifera* cv. Chardonnay vines planted E-W at 6-foot vine spacing and 9-foot row spacing. Each row contains sixteen panels, where a single panel has three or four vines depending on planting year. All vines are grafted on rootstock 3309 and were trained using a double mid-wire cordon method. Vines were spur-pruned in early spring of each year.

This investigation was conducted concurrently with annual fungicide efficacy trials in the chardonnay block. Each year, all fungicide treatments were replicated four times with each 3- or 4-vine panel representing a single replication. The trials were organized in a randomized complete block design (RCBD). Applications were made with a custom hooded boom sprayer attached to a 90-horsepower tractor, operating at 100 psi and delivering a volume of 50 gallons per acre (gpa) pre-bloom and 100 gpa post-bloom. Fungicides were applied once every seven to ten days from June to August each year, for seven total applications annually. Disease was caused by native pathogen populations with no manual inoculation. Weekly disease ratings began once the first visible symptoms appeared each year. In 2020 the trial area comprised the ten rows at the southern end of the vineyard, while in 2021 and 2022 we extended the study to all twenty rows. Treatment diagrams are provided in Supplementary Figure S1.

### Disease rating

We conducted the study over three growing seasons (June to August 2020, 2021, and 2022). Each year, a team of trained scouts recorded weekly GDM severity and incidence ratings. Scouts sampled twenty leaves per panel and recorded percent symptomatic leaf area for each leaf (Nutter and Schultz 1995). Tissue was considered symptomatic if it displayed characteristic yellow oil spots on the adaxial surface and sporulation on the abaxial leaf face (Figure 1) (Camargo et al. 2019). We averaged the twenty measurements to calculate severity for each panel. GDM severity ratings presented here are average percent symptomatic leaf area on a per-panel basis. We calculated GDM incidence as the number of symptomatic leaves divided by the total number of leaves sampled (Nutter and Schultz 1995). For example, if we sampled twenty leaves and four displayed GDM symptoms, the GDM incidence was recorded as 25%.

**Figure 1.**
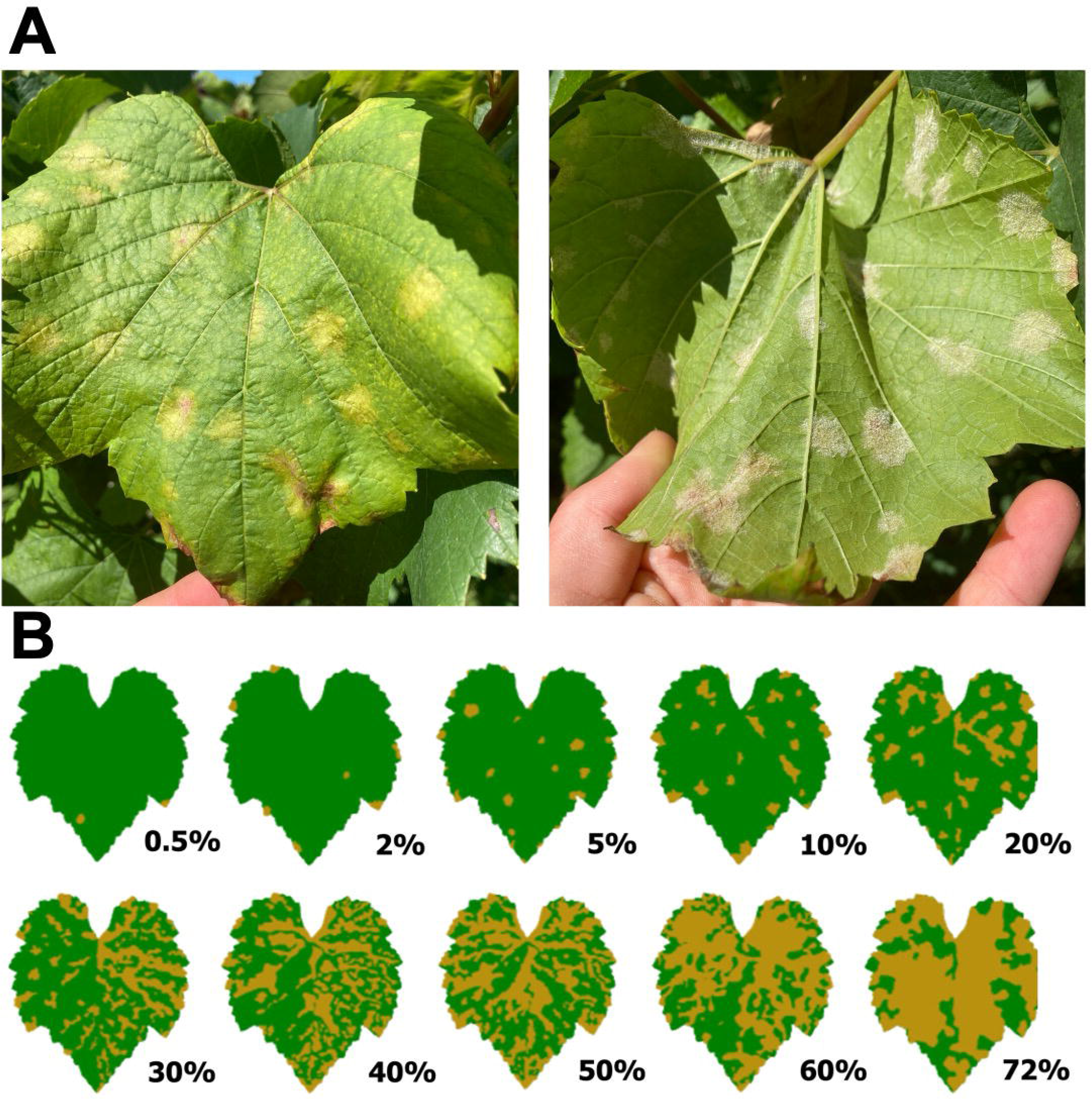
**A** Foliar downy mildew signs and symptoms: chlorotic lesions on the adaxial leaf face (left) with sporulation on the abaxial surface (right). **B** Standard area diagram used to rate downy mildew severity, adapted from Camargo et al. 2019.

### Satellite imagery

Each year PlanetScope and SkySat imagery was acquired between June 1 and August 31 over the study site. Table 1 summarizes the relevant technical specifications for both image types. We downloaded all imagery directly from Planet Explorer (https://www.planet.com/explorer). Both image product types include an orthorectified, atmospherically corrected GEOTIFF file with top-of-atmosphere (TOA) reflectance for each pixel. We selected images based on the following criteria:

- Acquired within 72 hours of GDM scouting
- Cloud-free
- Successful co-registration with a reference image

**Table 1.**
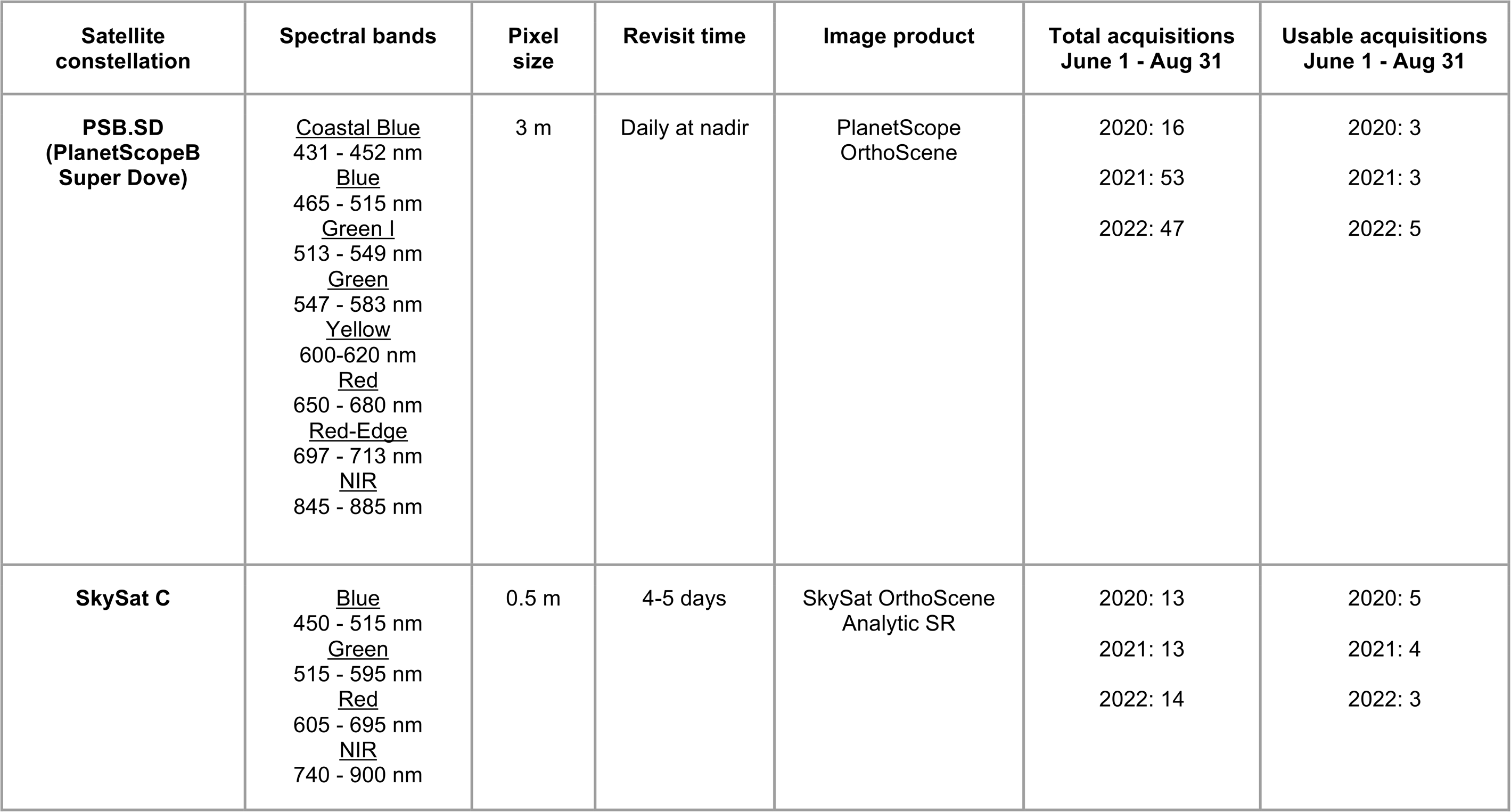
Satellite platform and image product specifications, adapted from “Planet Imagery Product Specifications”, May 2022.

We co-registered the selected images using the AROSICS (Automated and Robust Open-Source Coregistration Software for Multi-Sensor Satellite Data) python package (Scheffler et al. 2017). Specifically, we implemented the AROSICS global method, which calculates a single X/Y shift vector based on the displacement between pixels located at a single location in a reference and target image. The global method applies the same X/Y shift to all pixels in the target image without altering the pixel values, thus preserving all original TOA reflectance information. For a detailed explanation of the AROSICS package, refer to (Scheffler et al. 2017). After filtering by cloud cover and image date and performing co-registration, we had 3 and 5 (2020), 3 and 4 (2021), and 5 and 3 (2022) usable PlanetScope and SkySat images, respectively, across the three-year study period. No additional geometric or radiometric corrections were applied to the imagery beyond the commercial provider’s pre-processing steps (“Planet Imagery Product Specifications” 2022).

### Image labeling

For SkySat images, we generated a mask to isolate pixels within the footprint of each three- or four-vine panel. To ensure spatial accuracy of the mask we georeferenced all vineyard trellis posts using a Duro RTK GNSS receiver, which yields geolocation information at sub-centimeter accuracy (“Duro Product Summary” 2021). We used the georeferenced coordinates of each trellis post to generate a GEOJSON file containing the panel polygons between each pair of posts (Figure 2).

**Figure 2.**
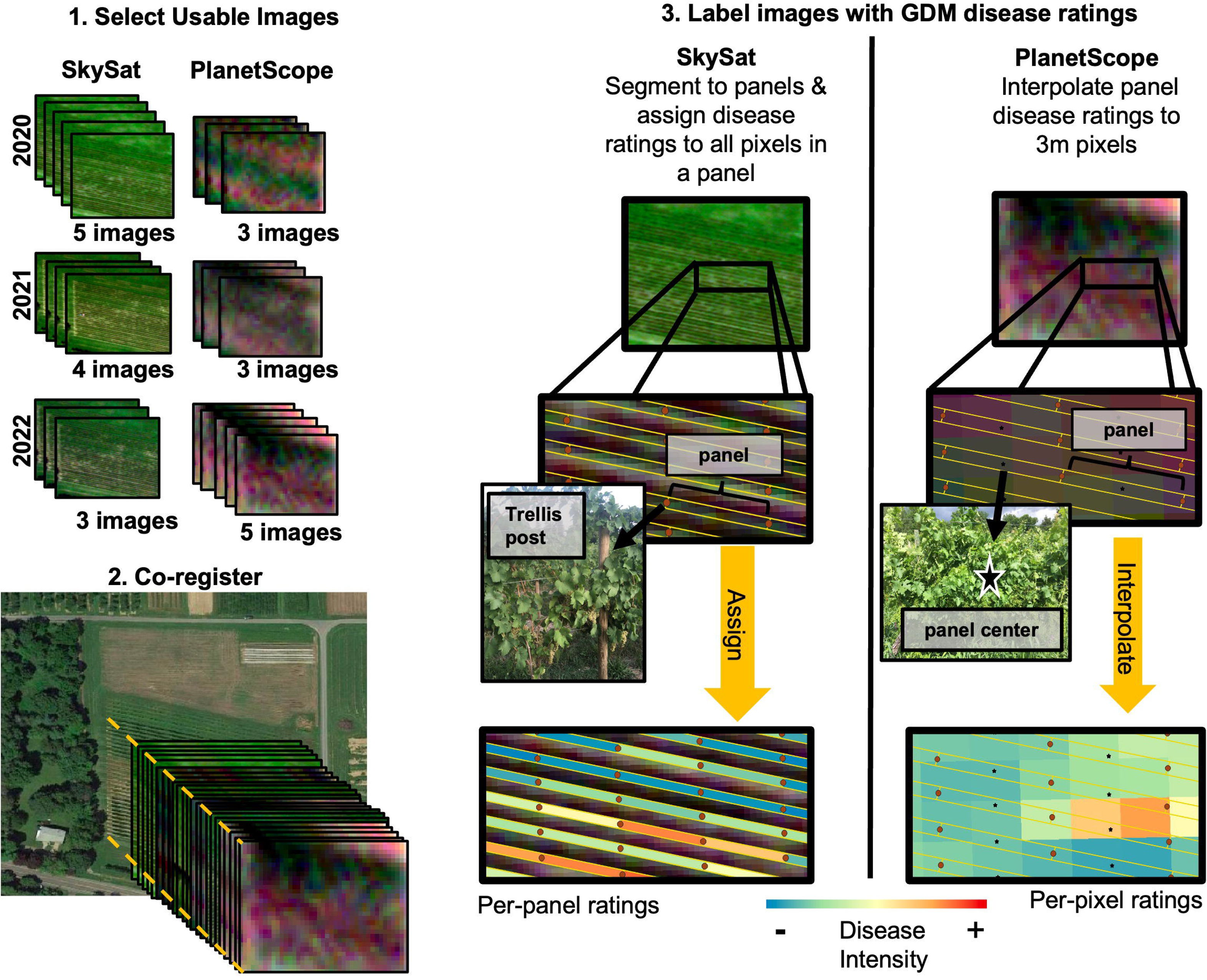
Image processing workflow

We assigned panels to “high” or “low” disease severity and incidence classes based on the scout rating taken closest to the image date. For disease severity, we assigned panels with greater than 10% average symptomatic leaf area to the high severity class. Panels with average symptomatic area less than or equal to 10% were assigned to the low severity class. Similarly, for disease incidence we assigned panels with greater than 25% incidence (more than five symptomatic leaves in a sample of twenty) to the high incidence class. We assigned panels with incidence less than or equal to 25% to the low incidence class.

Due to the lower spatial resolution of the PlanetScope images (3.9m ground sample distance, approximately 3m per pixel), each pixel overlaps multiple panels. This prevented extraction of reflectance data for individual panels in the PlanetScope images. Instead, we assigned the disease severity and incidence ratings for each panel to the X,Y coordinate at the centroid of the panel polygon. Using the centroid points, we interpolated disease severity and incidence over a 3 x 3 m grid to generate per-pixel severity and incidence ratings that match the spatial resolution of the PlanetScope images (Figure 2). We labeled pixels as high or low severity and incidence using the classification strategy described above.

### Top of atmosphere (TOA) reflectance and vegetation indices (VIs)

For both image datasets, we used TOA reflectance to calculate a series of multispectral vegetation indices. For SkySat we calculated the normalized difference vegetation index (NDVI), atmospherically resistant vegetation index (ARVI), modified soil adjusted vegetation index (MSAVI), enhanced vegetation index (EVI), and green-red vegetation index (GRVI). For eight-band PlanetScope images we also computed the photochemical reflectance index (PRI), the transformed chlorophyll absorption in reflectance index (TCARI) and the normalized difference red edge (NDRE). NDVI is the normalized difference ratio of near infrared to red reflectance and is the most widely used VI for remote sensing of terrestrial vegetation because it serves as a robust metric for differentiating vegetated from non-vegetated areas and is simple to calculate (Zeng et al. 2022; Huang et al. 2021). The ARVI, initially developed to improve upon NDVI measurements from MODIS-AVHRR data, uses blue reflectance to correct for atmospheric noise induced by variable aerosol content (Kaufman and Tanre 1992). The MSAVI is derived from the soil adjusted vegetation index (SAVI) and employs a dynamic correction factor to reduce noise from soil background reflectance (Qi et al. 1994). The enhanced vegetation index, EVI, corrects for both soil and atmospheric noise and, like ARVI, incorporates the blue band (Liu and Huete 1995). The GRVI is the normalized difference ratio of green to red reflectance and has been shown to correlate closely with green biomass, total chlorophyll, and gross primary productivity (Tucker 1979; Yin et al. 2022). The additional indices calculated for PlanetScope imagery take advantage of narrower spectral regions. Gamon et al. found that PRI, the normalized difference ratio of reflectance at 539 and 550 nm, correlates closely with xanthophyll cycle pigment activity and can serve as a proxy for absorbed photosynthetically active radiation (Gamon, Peñuelas, and Field 1992). Similarly, TCARI and NDRE take advantage of higher spectral resolution in the red to red-edge region to approximate chlorophyll content (Sims and Gamon 2002; Haboudane et al. 2002). We selected VIs based on previous work demonstrating that these indices are robust indicators of plant health, especially when soil and shadow contribute significantly to the background reflectance (Zeng et al. 2022; Giovos et al. 2021).

For SkySat datasets, we computed VIs for each panel by averaging the VIs of all pixels within the panel. VIs for PlanetScope are per-pixel. To identify VIs that can serve as indicators of GDM damage, we computed the Spearman’s rank correlation between each VI, GDM severity, and GDM incidence. Spearman’s rank is a nonparametric test of the linear correlation between two variables that is appropriate when values of at least one variable are not normally distributed and/or when samples being compared are of different sizes, as is the case with the GDM incidence and severity ratings (see Supplementary Figure S2) (Daniel 1990). We repeated this test for SkySat and PlanetScope datasets separately. For each dataset, we identified the three VIs with the largest negative correlation coefficients across the three study years, indicating the strongest inverse relationship to disease incidence or severity. This is based on previous work demonstrating that VIs indicate vegetation health status (Glenn et al. 2008; Zeng et al. 2022; Yao et al. 2012; Haboudane et al. 2002; Barreto et al. 2023; di Gennaro et al. 2016; Calderón, Navas-Cortés, and Zarco-Tejada 2015), such that relatively lower VI values can signal plant stress. Table 2 and Table 3 list VIs with associated Spearman correlation coefficients and p-values for SkySat and PlanetScope, respectively.

**Table 2.** Spearman’s correlations between GDM severity and incidence and SkySat vegetation indices (VIs), 2020 - 2022. ^a^ Bold entries indicate the VIs with the largest negative correlation coefficients. ^b^ Asterisks *, **, and *** indicate significance at the P ≤ 0.05, 0.01, and 0.001 levels. Bold values indicate the correlation coefficients for the three VIs with largest negative Spearman’s r for incidence or severity.

| VI<sup>a</sup> | Formula | Citation | Spearman's r<sup>b</sup> |  |  |  |  |  |
| --- | --- | --- | --- | --- | --- | --- | --- | --- |
|  |  |  | 2020 |  | 2021 |  | 2022 |  |
|  |  |  | Incidence | Severity | Incidence | Severity | Incidence | Severity |
| NDVI | $(NIR - RED) / (NIR + RED)$ | Rouse et al. (1974) | 0.62*** | 0.65*** | -0.17*** | -0.12*** | 0.64*** | 0.06 |
| EVI | $(2.5 * (NIR - RED)) / (NIR + 6 * RED - 7.5 * BLUE + 1)$ | Huete et al. (1994); Liu and Huete (1995) | 0.57*** | 0.59*** | **-0.22\*\*\*** | -0.18*** | 0.60*** | 0.08* |
| MSAVI | $((2 * (NIR) + 1 - \sqrt{((2 * NIR + 1) * 2 - 8 * (NIR - RED)))}) / 2)$ | Qi et al. (1994) | 0.57*** | 0.59*** | -0.14*** | -0.12*** | 0.39*** | 0.27*** |
| ARVI | $(NIR - (2 * RED - BLUE)) / (NIR + (2 * RED - BLUE))$ | Kaufman and Tanré (1992) | 0.61*** | 0.63*** | **-0.35\*\*\*** | -0.30*** | 0.60*** | -0.03 |
| GRVI | $(GREEN - RED) / (GREEN + RED)$ | Tucker (1979) | 0.61*** | 0.62*** | **-0.33\*\*\*** | -0.28*** | 0.38*** | -0.19*** |

**Table 3.** Spearman’s correlations between GDM severity and incidence and PlanetScope vegetation indices (VIs), 2020 - 2022. ^a^ Asterisks *, **, and *** indicate significance at the P ≤ 0.05, 0.01, and 0.001 levels.

| VI | Formula | Citation | Spearman's $r^a$ | | | | | |
| --- | --- | --- | --- | --- | --- | --- | --- | --- |
|  |  |  | 2020 |  | 2021 |  | 2022 |  |
|  |  |  | Incidence | Severity | Incidence | Severity | Incidence | Severity |
| **NDVI** | $(NIR - RED) / (NIR + RED)$ | Rouse Jr. et al. 1974 | -0.29*** | -0.15** | **-0.39\*\*\*** | -0.30*** | 0.07*** | 0.08*** |
| EVI | $(2.5 * (NIR - RED)) / (NIR + 6 * RED - 7.5 * BLUE + 1)$ | Huete, Justice, and Liu 1994; Liu and Huete 1995 | -0.31*** | -0.17** | -0.35*** | -0.28*** | 0.07*** | 0.07*** |
| MSAVI | $((2 * (NIR) + 1 - \sqrt{((2 * NIR + 1) * 2 - 8 * (NIR - RED)))}) / 2)$ | Qi et al. 1994 | -0.30*** | -0.22*** | -0.33*** | -0.28*** | 0.07*** | 0.08*** |
| ARVI | $(NIR - (2 * RED - BLUE)) / (NIR + (2 * RED - BLUE))$ | Kaufman and Tanre 1992 | -0.28*** | -0.11 | -0.29*** | -0.25*** | 0.06*** | 0.07*** |
| GRVI | $(GREEN - RED) / (GREEN + RED)$ | Tucker 1979 | -0.34*** | -0.18** | -0.34*** | -0.33*** | 0.02 | 0.08*** |
| **PRI** | $(GREEN - GREEN1) / (GREEN + GREEN1)$ | Gamon, Peñuelas, and Field 1992 | -0.35*** | -0.28*** | **-0.41\*\*\*** | -0.26*** | -0.02 | 0.02 |
| **TCARI** | $(3 * (REDE - RED) - 0.2 * (REDE - GREEN)) * (REDE / RED)$ | (Haboudane et al. 2002) | -0.28*** | -0.16** | **-0.37\*\*\*** | -0.31*** | 0.03 | 0.02 |
| NDRE | $(NIR - REDE) / (NIR + REDE)$ | Sims and Gamon 2002 | -0.12* | -0.05 | -0.24*** | -0.21*** | 0.09*** | 0.14*** |

### VIs by image date

To determine how early in the growing season we could detect disease-induced differences in VIs, we calculated the Mann-Whitney U statistic and p-value for high and low intensity classes on each image date. The Mann-Whitney U is a nonparametric statistic used to determine whether two samples come from the same population. Like the Spearman’s rank coefficient, the U statistic is calculated by first ranking samples and then computing the U statistic based on sample rank rather than the raw value (Mann and Whitney 1947). We repeated this process on each image date, with both SkySat and PlanetScope imagery (Figure 3 and Figure 4).

**Figure 3.**
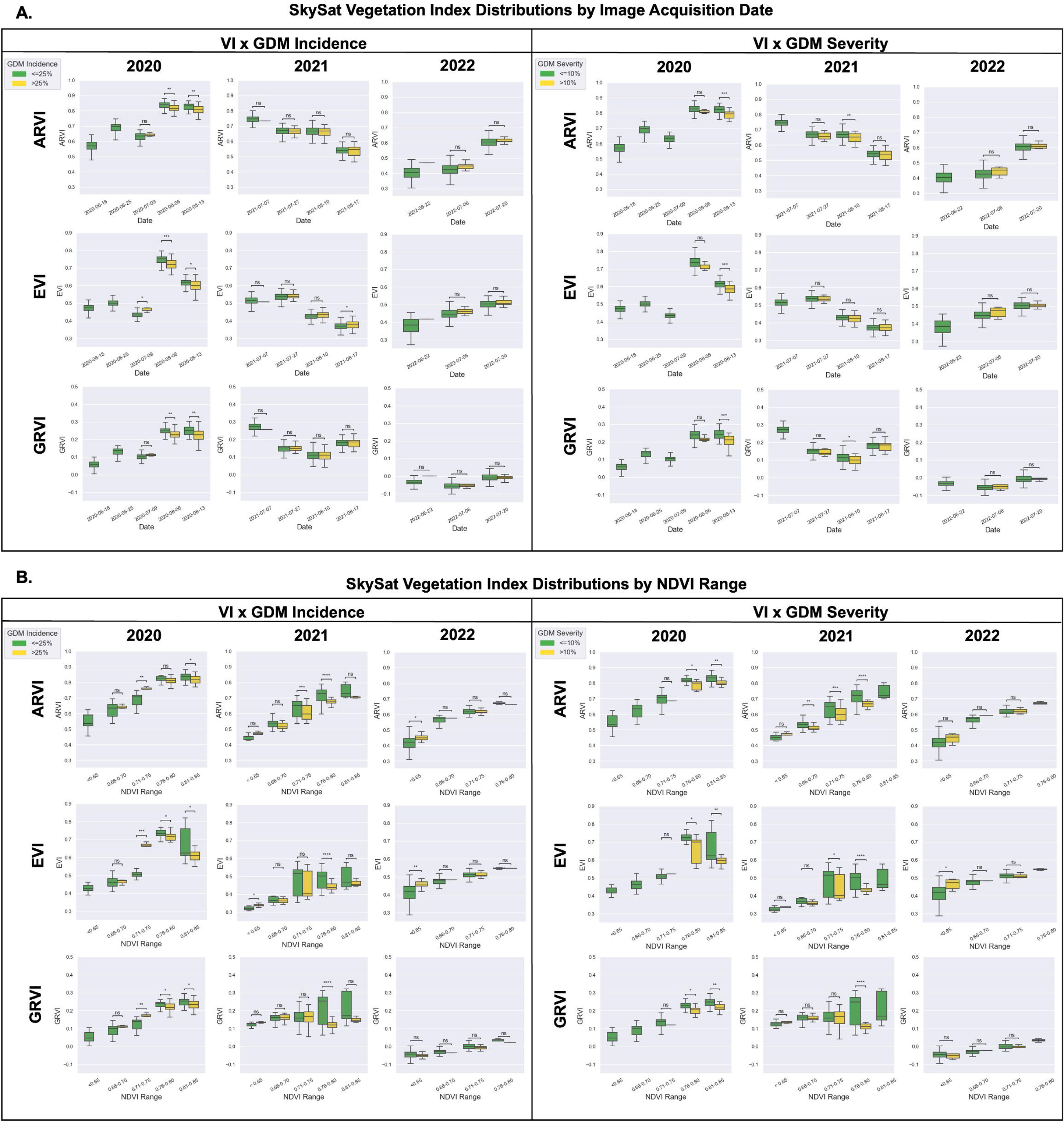
SkySat ARVI, EVI and GRVI values for grapevine panels exhibiting low and high GDM incidence and severity. Panels are aggregated by image acquisition date (**A**) and by NDVI range (**B**) in 2020, 2021, and 2022. Between-group differences were evaluated by the Mann-Whitney test. Asterisks *, **, ***, and **** indicate significant differences between severity and incidence groups at the P ≤ 0.05, 0.01, 0.001, and 0.0001 levels.

**Figure 4.**
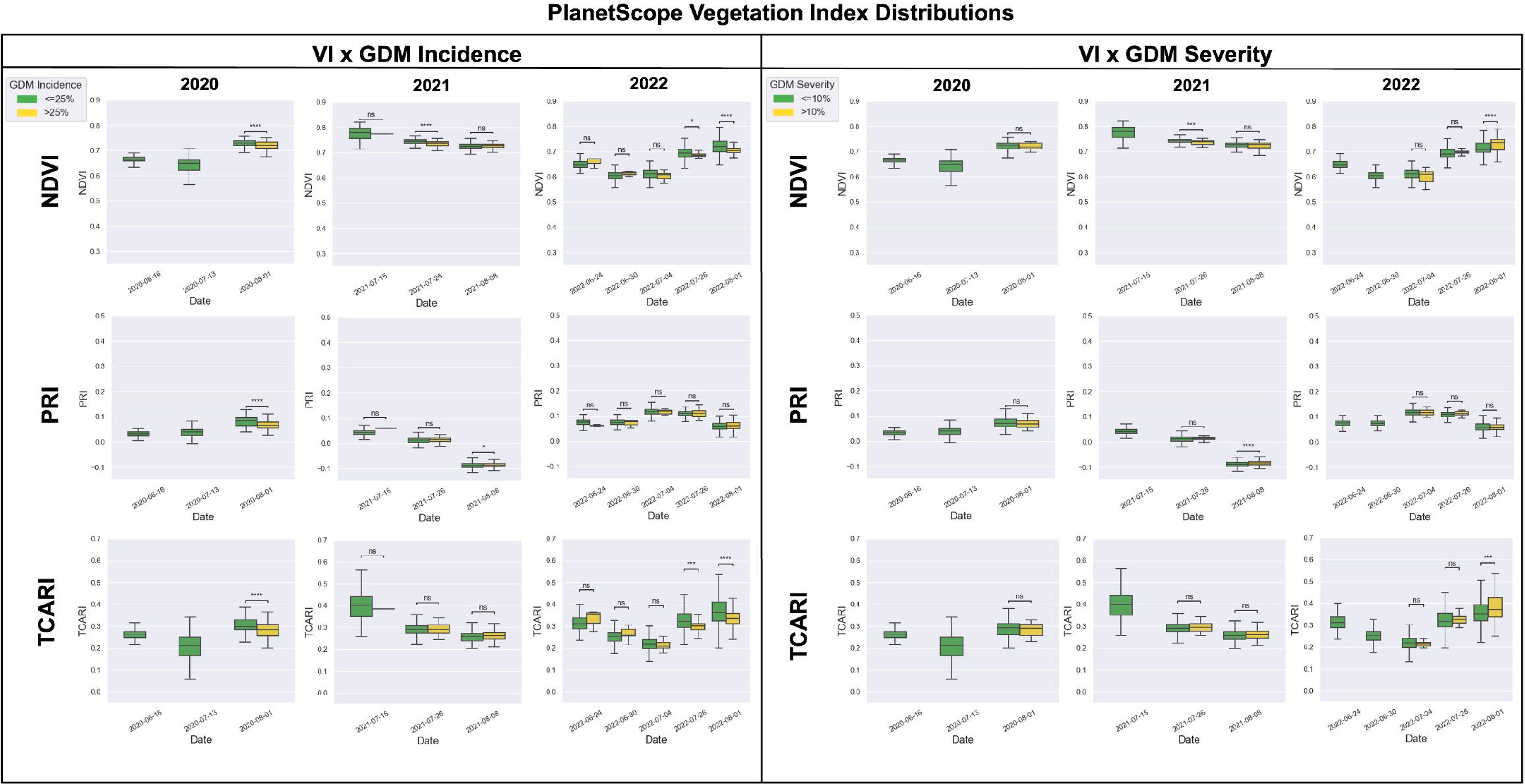
NDVI, PRI and TCARI distributions for PlanetScope pixels assigned low and high GDM incidence and severity based on interpolated disease ratings. Between-group differences were evaluated by the Mann-Whitney test. Asterisks *, **, ***, and **** indicate significant differences between groups at the P ≤ 0.05, 0.01, 0.001, and 0.0001 levels.

### SkySat VIs by NDVI

To account for differences in canopy size in the higher resolution SkySat imagery, we aggregated panels by NDVI and compared VI values of high and low intensity (both incidence and severity) GDM classes within NDVI brackets. This follows work by Calderón et al. demonstrating that aggregating crop pixels by NDVI ranges enhanced detection of downy mildew in opium poppy, with asymptomatic and symptomatic plants having significantly different ratios of green to red reflectance at high NDVI steps (Calderón et al. 2014). We performed our analysis by first pooling all data across one season and then grouping grapevine panels by NDVI range. Thus, a single NDVI bracket can include panels from images acquired on different dates within the same year. NDVI is used here as a proxy for total green leaf area and the relative size of the grapevine canopy within a panel.

### Random Forest Classifiers for SkySat and PlanetScope imagery

To evaluate the potential for automated detection of areas with high GDM damage, we trained random forest classifiers using different combinations of spectral bands and spectral bands plus vegetation indices as input features. Random forest classification is a widely used method in agricultural remote sensing and disease detection (Vitrack-Tamam et al. 2020; Iordache et al. 2020; Heim et al. 2019; Hassanzadeh et al. 2021; Guo et al. 2022; Galvan et al. 2023; Gold et al. 2020). For each combination of year (2020, 2021, 2022, and pooled data 2020-22), satellite platform (SkySat and PlanetScope), and disease metric (severity and incidence) we built one model using only the spectral band reflectance (SB) and one model using spectral bands and VIs (SB + VI) to evaluate whether including VIs improved model accuracy. This yielded a total of 4 years x 2 platforms x 2 metrics x 2 input feature sets = 32 random forest models (Table 4 and 5).

**Table 4.** Random forest model results using SkySat spectral bands (SB) or spectral bands and vegetation indices (SB +VI) to classify panels by GDM severity or incidence. Vegetation indices are NDVI, EVI, MSAVI, ARVI, GRVI. Bolded values are accuracy scores for the best performing model in each category.

| SkySat Random Forest GDM Severity |  |  |  |  |  |  |  |  |
| --- | --- | --- | --- | --- | --- | --- | --- | --- |
|  | 2020 |  | 2021 |  | 2022 |  | All years |  |
|  | SB | SB + VI | SB | SB + VI | SB | SB + VI | SB | SB + VI |
| Precision high severity | 0.75 | 0.74 | 0.73 | 0.73 | 0.59 | 0.41 | 0.75 | 0.74 |
| Precision low severity | 0.74 | 0.78 | 0.81 | 0.80 | 0.64 | 0.45 | 0.77 | 0.78 |
| Recall high severity | 0.72 | 0.78 | 0.83 | 0.82 | 0.65 | 0.47 | 0.77 | 0.79 |
| Recall low severity | 0.73 | 0.71 | 0.69 | 0.69 | 0.56 | 0.46 | 0.73 | 0.72 |
| F1 high severity | 0.72 | 0.75 | 0.77 | 0.77 | 0.60 | 0.42 | 0.76 | 0.76 |
| F1 low severity | 0.72 | 0.73 | 0.74 | 0.74 | 0.56 | 0.43 | 0.75 | 0.74 |
| Overall accuracy | 0.73 | 0.74 | 0.76 | 0.76 | 0.61 | 0.47 | 0.75 | 0.76 |
| SkySat Random Forest GDM Incidence |  |  |  |  |  |  |  |  |
|  | 2020 |  | 2021 |  | 2022 |  | All years |  |
|  | SB | SB + VI | SB | SB + VI | SB | SB + VI | SB | SB + VI |
| Precision high incidence | 0.85 | 0.84 | 0.72 | 0.72 | 0.61 | 0.59 | 0.78 | 0.77 |
| Precision low incidence | 0.91 | 0.91 | 0.76 | 0.75 | 0.66 | 0.61 | 0.81 | 0.80 |
| Recall high incidence | 0.92 | 0.92 | 0.78 | 0.77 | 0.71 | 0.65 | 0.82 | 0.80 |
| Recall low incidence | 0.84 | 0.83 | 0.70 | 0.69 | 0.54 | 0.54 | 0.77 | 0.76 |
| F1 high incidence | 0.88 | 0.88 | 0.75 | 0.74 | 0.65 | 0.61 | 0.80 | 0.79 |
| F1 low incidence | 0.87 | 0.87 | 0.73 | 0.72 | 0.58 | 0.56 | 0.79 | 0.78 |
| Overall accuracy | 0.88 | 0.87 | 0.74 | 0.73 | 0.62 | 0.60 | 0.79 | 0.78 |

**Table 5.** PlanetScope random forest model results using spectral bands as input features (SB) or spectral bands and vegetation indices (SB +VI) to classify GDM severity and incidence. Vegetation indices are NDVI, EVI, MSAVI, ARVI, GRVI, PRI, TCARI, and NDRE. Bolded values are accuracy scores for the best performing model in each category.

| PlanetScope Random Forest GDM Severity |  |  |  |  |  |  |  |  |
| --- | --- | --- | --- | --- | --- | --- | --- | --- |
|  | 2020 |  | 2021 |  | 2022 |  | All years |  |
|  | SB | SB + VI | SB | SB + VI | SB | SB + VI | SB | SB + VI |
| Precision high severity | 0.85 | 0.84 | 0.82 | 0.80 | 0.81 | 0.83 | 0.81 | 0.80 |
| Precision low severity | 0.99 | 0.97 | 0.89 | 0.87 | 0.82 | 0.84 | 0.85 | 0.87 |
| Recall high severity | 0.99 | 0.97 | 0.90 | 0.88 | 0.82 | 0.84 | 0.86 | 0.88 |
| Recall low severity | 0.80 | 0.80 | 0.79 | 0.78 | 0.80 | 0.82 | 0.80 | 0.78 |
| F1 high severity | 0.91 | 0.89 | 0.85 | 0.84 | 0.81 | 0.83 | 0.83 | 0.84 |
| F1 low severity | 0.88 | 0.87 | 0.84 | 0.82 | 0.81 | 0.83 | 0.82 | 0.82 |
| Overall accuracy | **0.90** | 0.88 | 0.85 | 0.83 | 0.81 | 0.83 | 0.83 | 0.83 |
| PlanetScope Random Forest GDM Incidence |  |  |  |  |  |  |  |  |
|  | 2020 |  | 2021 |  | 2022 |  | All years |  |
|  | SB | SB + VI | SB | SB + VI | SB | SB + VI | SB | SB + VI |
| Precision high incidence | 0.91 | 0.91 | 0.79 | 0.79 | 0.81 | 0.79 | 0.84 | 0.85 |
| Precision low incidence | 0.99 | 0.99 | 0.84 | 0.82 | 0.79 | 0.78 | 0.86 | 0.86 |
| Recall high incidence | 0.99 | 0.99 | 0.86 | 0.83 | 0.78 | 0.77 | 0.87 | 0.86 |
| Recall low incidence | 0.90 | 0.90 | 0.77 | 0.78 | 0.82 | 0.80 | 0.83 | 0.85 |
| F1 high incidence | 0.95 | 0.95 | 0.82 | 0.81 | 0.79 | 0.78 | 0.85 | 0.86 |
| F1 low incidence | 0.94 | 0.94 | 0.80 | 0.80 | 0.80 | 0.79 | 0.85 | 0.85 |
| Overall accuracy | **0.94** | 0.94 | 0.81 | 0.80 | 0.80 | 0.81 | 0.85 | 0.85 |

The GDM severity and incidence data is imbalanced across all years (see Supplementary Figure S3). To address class imbalance, we used the imbalanced-learn library to randomly under sample the low damage classes (both severity and incidence) to match the size of the high damage classes in each year (Lemaitre, Nogueira, and Christos 2017). We split the balanced datasets in a 70/30 train/test ratio and normalized the feature values to a standard range from 0-1 with a minimum/maximum scaler. After splitting and normalization, we fit a baseline random forest model with each dataset and computed accuracy, precision, recall, and F1 scores for both the high and low classes. Next, we tuned hyperparameters of each model using the scikit-learn RandomSearchCV and GridSearchCV modules (Pedregosa et al. 2011). The tuned hyperparameters were:

- n_estimators: the number of trees in each random forest
- max_features: the number of features considered at each split
- max_depth: the maximum number of splits per tree
- min_samples_split: the minimum number of samples that can trigger a split
- min_smaples_leaf: the minimum number of samples at each leaf in the tree
- bootstrap: whether to sample with replacement from the original dataset for each tree

We fit the baseline and tuned random forest models with the normalized, scaled training data and applied fitted models to the test set. We initiated each model for 100 random train/test splits and calculated average accuracy, precision, recall, and F1 scores. We compared metrics between the baseline and tuned models to select the best performing model for each dataset. We calculated the permutation feature importance to determine which features were most important for classification in all final models. Random forest results are presented in Table 4 (SkySat) and Table 5 (PlanetScope). All models were implemented using the scikit-learn python library (Pedregosa et al. 2011).

## III. Results

### Disease development

We evaluated GDM intensity by disease incidence and disease severity. GDM incidence reached 100% in at least one panel in all three years, while maximum GDM severity on the final day of scouting varied from 57 to 82% (Supplementary Table S1). However, most panels maintained low levels of disease intensity due to regular fungicide applications. Average GDM severity and incidence on the final day of scouting ranged from 3 to 14% and 22 to 81%, respectively (Supplementary Table S1).

Considering only dates with concurrent scouting and satellite image acquisitions, panels in the high incidence class (GDM incidence > 25%) were not observed until 09-July in 2020, 15-July in 2021, and 29-June in 2022 (Figure S3). Across all years, panels exceeded the 25% incidence threshold earlier than the 10% mean severity threshold. In 2020, panels with mean GDM severity exceeding 10% were not observed until 30 July (Figure S3). PlanetScope imagery provided better temporal coverage of late-stage GDM than SkySat in 2022. On the last SkySat acquisition date in 2022 (2022-07-20), the ratios of low to high incidence and severity panels were 17.5 to 1 and 59 to 1, respectively. In the same year, the final PlanetScope acquisition date (2022-08-02) captured an increase in both incidence and severity with low to high ratios of 4.6 to 1 and 3.7 to 1, respectively.

### SkySat VI distributions by image date

SkySat VIs differed significantly (P<0.05) between disease incidence classes more often than between disease severity classes (Figure 3A) when panel ratings were aggregated by image date. For disease severity and incidence, the median VI values of the high intensity classes exceeded those of low intensity classes on multiple dates, though these differences were only significant between the high and low incidence groups on 2021-08-17 for EVI (Figure 3A) when aggregated by image date.

On the two latest SkySat image acquisition dates in 2020 the number of high incidence panels (GDM incidence >25%) exceeded the number of low incidence panels by more than ten percent, while the number of high severity panels was never greater than the number of low severity panels in any of the three years. On the dates when the high incidence panel count exceeded the low incidence panel count (2020-08-06 and 2020-08-13), significant differences between incidence classes were observed in all three VIs (Figure 3A). The only other date when we observed significant differences between SkySat VIs of distinct GDM incidence classes was 2021-08-17, when the number of high incidence panels approached thirty percent the number of low incidence panels. However, on 2021-08-17 the direction of the difference changed, with median high-incidence VIs exceeding median low-incidence VIs (Figure 3A).

High severity panels had significantly lower EVI, ARVI and GRVI than low severity panels on the final image date in 2020, when the number of high severity panels reached a maximum, while no significant differences between severity classes were observed in any SkySat VIs on the final image dates in 2021 or 2022. Significantly lower ARVI and GRVI were observed in high severity panels on the second-to-last image date in 2021 (2021-08-10), but differences in EVI were not significant at the P<0.05 level (Figure 3A).

### SkySat VI distributions by canopy size

As observed with VIs grouped by image date, SkySat VIs differed significantly (P<0.05) between GDM incidence classes more often than between severity classes when panels are grouped by NDVI ranges as a proxy for canopy size (Figure 3B). In 2020 the lower NDVI brackets (0.66-0.70, and 0.71-0.75) corresponded to higher EVI, GRVI, and ARVI for panels with >25% GDM incidence, with lower VI values observed in healthier panels. However, these differences only reached significance at the P<0.05 level in 2020 for the 0.71-0.75 NDVI bracket. The same pattern occurred in 2021 and 2022, with high incidence panels in the ≤0.65 NDVI bracket having larger values for all three VIs than low incidence panels. However, as we observed in 2020 most of these differences were not significant at the P<0.05 level (Figure 3B). EVI and GRVI were lower for high incidence panels in the 0.76-0.80 NDVI bracket in 2020 and 2021, and in the 0.81-0.85 bracket in 2020 (P<0.05). No significant differences between GDM incidence groups were observed in ranges above 0.65 for any VIs in 2022.

We observed the same result in 2022 when using GDM severity as the intensity metric, with high severity (>10% symptomatic leaf area) panels exhibiting significantly greater (P<0.05) median EVI within the <0.65 bracket but no other significant differences in VIs between severity classes for later image dates. In 2020 and 2021 the only significant differences between severity classes indicate smaller VI values for high severity panels and larger values for healthy panels. These differences are significant across all three VIs for panels with NDVI greater than 0.75 in 2020 and with NDVI between 0.76 and 0.80 in 2021 (Figure 3B).

### PlanetScope VI distributions

As we observed with the SkySat dataset, GDM incidence was a better metric in terms of identifying significant differences between high and low disease intensity classes. Values of at least one VI for pixels of high and low incidence classes were significantly different at the P<0.05 level on dates in 2020, 2021, and 2022. Apart from the median PRI values on 08-08-2021, all other significant between-class differences indicated larger VI values for healthy pixels than high incidence pixels. The only dates on which PlanetScope VIs differed significantly at the P<0.05 level between disease classes for both incidence and severity were 2022-08-01 (NDVI and TCARI) and 2021-07-26 (NDVI) (Figure 4).

Severity classes did not differ significantly in any VIs in 2020, while in 2021 significant differences were observed in NDVI and PRI on 26-July and 08-Aug, respectively. However, while median NDVI was lower for high severity pixels than low severity pixels, median PRI was larger. In 2022 NDVI and TCARI values were significantly larger for high severity pixels on the final image date, 2022-08-01 (Figure 4).

PlanetScope NDVI proved the best VI for distinguishing between incidence classes in all three years, with significantly higher values observed in the low incidence pixels on August image dates in 2020 and 2022, and on 26-July in 2021 (P<0.05). TCARI performed similarly to NDVI for distinguishing between incidence groups in 2020 and 2022, with significant differences observed in August images in both years and on 26- July in 2022. PRI values were significantly higher (P<0.0001) for low incidence than high incidence pixels in August 2020, but not in 2021 or 2022 (Figure 4).

As in the SkySat dataset, the August 2020 PlanetScope image revealed significantly lower values of all three VIs for high severity and incidence classes as compared to low severity and incidence groups. The August 2020 acquisitions correspond to the highest observed incidence and severity in 2020, with high incidence panels outnumbering low incidence panels and 25% of all panels falling in the high GDM severity group (Supplementary Figure S4).

### Random forest model performance

#### SkySat

Random forest classification models performed variably across years and input feature sets. Of the GDM severity classifiers, the model trained on spectral bands (SB) of 2021 images performed best, with an overall accuracy of 0.76 (Table 4). Across all severity models except for 2020 SB, precision was higher for low severity panels than high severity panels. The models built using SB and VIs had higher overall accuracy and F1 scores than the SB-only models in 2020 and in the pooled dataset but performed equivalently in 2021. The worst performing model is 2022 SB+VI with an overall accuracy score of 0.47. This model was trained on a SkySat dataset of only three images, with the latest acquisition on 26-July.

The models trained on SB and SB + VI data had nearly identical metrics for the 2020 GDM incidence classification task, with accuracy scores and high-incidence F1 of 0.88 and 0.88 and 0.87 and 0.88, respectively. The 2020 SB-only classifier had the highest overall accuracy of all the GDM incidence models and was the best performing GDM intensity classifier (considering both incidence and severity). The next-best performing model was the classifier trained on the 2020 SB+VI data. In 2021 the two incidence classifiers had similar precision, recall, accuracy, and F1 scores. The 2022 models had the lowest accuracy and F1 scores of the GDM incidence classifiers, though the 2022 SB + VI model performed better at incidence classification than the 2022 SB + VI model at severity classification (overall accuracies of 0.60 and 0.47, respectively). Although SB and SB+VI GDM incidence classifiers performed very similarly within each year, the SB-only models had slightly higher accuracy and F1 scores than their SB+VI counterparts. As observed in the severity classifiers, precision was higher for low incidence than high incidence classes across all input feature types and years (Table 4).

Finally, we tested the robustness of the model with the smallest input feature set (SB) trained on the greatest number of images (2020-2022). To see how a random forest classifier performs when GDM incidence is at a maximum, we applied the SkySat model trained on the SB 2020-2022 dataset to classify panels in an image acquired on August 16, 2021. We observed a decrease in accuracy, precision, and recall with respect to the randomly selected test set (Table 4 and Figure 5. We observed the same result when we applied the PlanetScope model trained on the three-year SB dataset to an image from August 01, 2022 (Figure 5). We selected these image dates as test cases because the proportion of high incidence panels (SkySat) and pixels (PlanetScope) had reached a maximum in both years (Figure S3).

**Figure 5.**
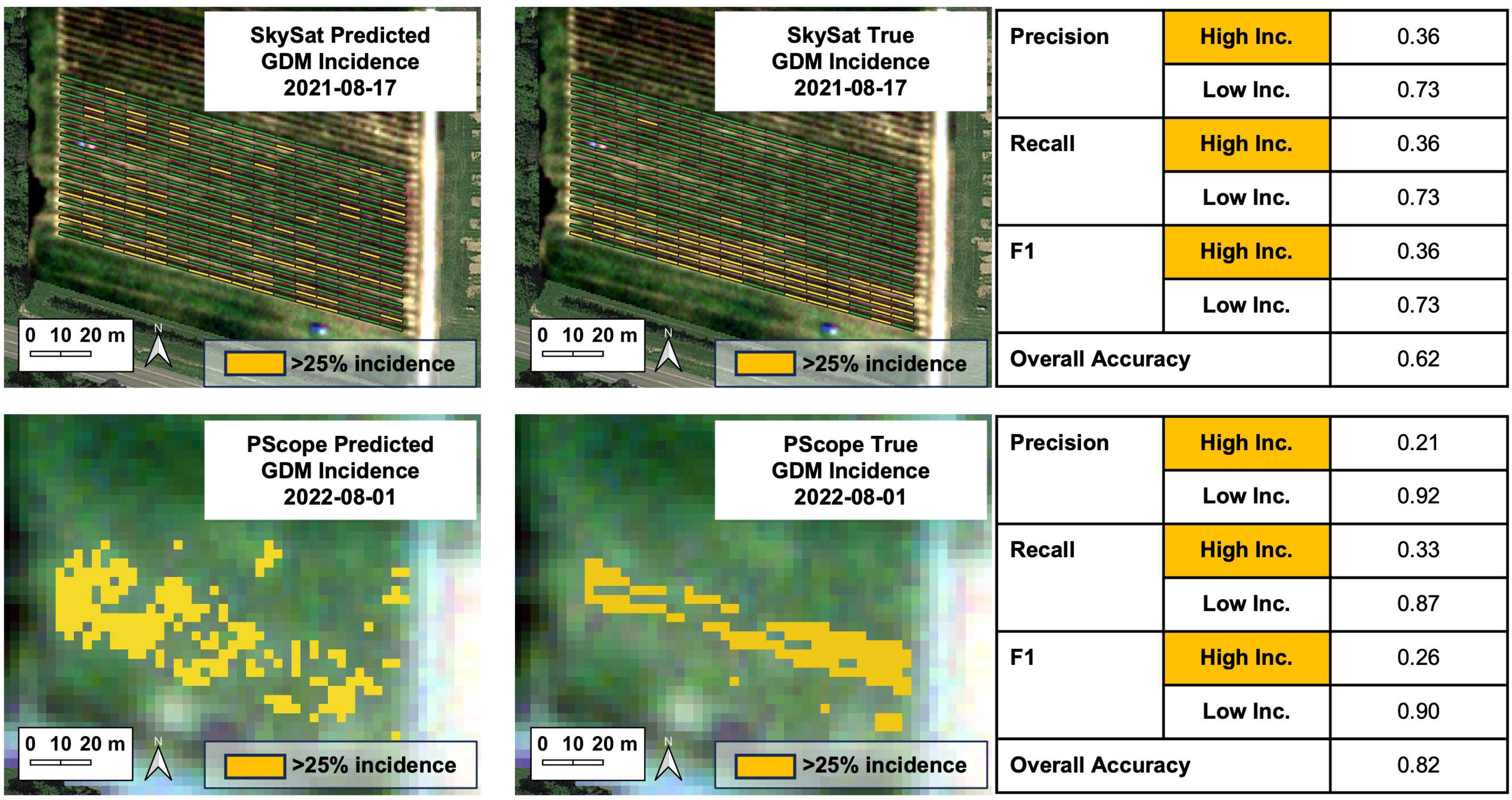
True GDM incidence ratings (right) and model predictions (left) of SkySat and PlanetScope random forest classifiers trained on the complete 2020-2022 spectral band dataset for each image type. Results shown are for a SkySat image acquired on August 17, 2021 and a PlanetScope image acquired on August 01, 2022.

#### PlanetScope

In contrast to the SkySat classifiers, the PlanetScope random forest models were trained and validated on the interpolated incidence and severity data with each pixel assigned to an incidence or severity class based on a GDM rating calculated based on the pixel’s distance to the center of surrounding panels and the incidence/severity recorded for these panels (Figure 2). The models trained on PlanetScope reflectance data all achieved accuracy scores of 0.80 or higher for classifying pixels as high or low incidence/severity (Table 5). For both incidence and severity, the most accurate classifiers were those trained on the 2020 dataset and the least accurate were those trained on 2022 data. However, the 2022 classifiers still performed well with OA of 0.81 (SB) and 0.80 (SB+VI) for incidence and 0.81 (SB) and 0.83 (SB+VI) for severity. For the classifiers trained on the complete three-year dataset, incidence was slightly better classified than severity (OA of 0.85 and 0.83, respectively).

As with SkySat, we observed only slight differences in performance metrics between classifiers trained on the SB and SB+VI input features. Unlike the SkySat data, classifier performance did not decrease substantially in 2022. In 2022 there were five total PlanetScope acquisitions, with the latest image acquired on 2022-08-01 (Supplementary Figure S3). Nevertheless, we observed a 3% decrease in overall accuracy when we applied the all-years model to classify pixels by incidence in the 2023-08-01 image (Figure 5). Precision and recall for the high incidence class dropped substantially, from 0.84 and 0.87 on the random 30% test set to 0.21 and 0.33, respectively, for the August 01 image (Table 4 and Figure 5).

The range of OA values across all years and input feature types was smaller for the PlanetScope data (OA from 0.80 to 0.95), than for SkySat (OA from 0.47 to 0.88). Classifier precision was higher for low severity and incidence classes than high classes across all models apart from the SB+VI incidence classifier in 2021 and both incidence classifiers in 2022.

## IV. Discussion

We sought to determine whether commercially available, high spatial resolution satellite imagery could be used to track GDM progression with multispectral plant health indicators and machine learning across multiple growing seasons. We investigated this capacity in a Chardonnay vineyard where variable fungicide applications established an artificial disease gradient. Overall, we found that both 50 cm and 3 m data streams were sufficient to detect GDM outbreaks above the economic injury threshold (Carlson and Headley 1987). We identified significant differences in vegetation indices between GDM diseased vs. non/minimally diseased grapevine late in the growing season, once average severity and incidence exceeded 10% and 25% in at least 7% and 16% of panels, respectively (Figures 3 and 4, Supplementary Figure S3). However, we found that disease surveillance capacity was limited by inconsistent availability of cloud-free, accurately orthorectified imagery, and thus differed across each of the three years in our study. In years when sufficient imagery was available during late infection stages, we observed consistent significant differences (P <0.05) between high and low severity/incidence classes in EVI, ARVI, and GRVI derived from SkySat images, and in PlanetScope NDVI. Random forest classifiers trained on individual spectral bands and VIs also performed better on datasets from years with a greater proportion of high severity/incidence samples, with the best overall accuracy for GDM detection in 2020 (with both SkySat [50 cm] and PlanetScope [3 m]) when high-incidence panels outnumbered low-incidence panels on the final image dates (Tables 4 and 5, Supplementary Figure S3). Thus, while we establish the utility and innate capacity of this imagery to achieve GDM detection, we identify operational barriers to enacting the overarching goal of using these data streams for disease surveillance.

Our findings align with other investigations using commercially available high resolution multispectral satellite imagery for disease detection. Guo et al. could distinguish healthy (< 1% symptomatic leaf area) from diseased areca nut canopies with PlanetScope imagery using a random forest model with overall accuracy of 0.88 (Guo et al. 2022). However, unlike a vineyard, the areca nut cropping system has a closed canopy and sample locations were 3×3 meter plots of 100% target foliage, with no soil or non-target vegetation contributing to the reflectance signal. Raza et al. found that a random forest model trained on PlanetScope spectral bands and NDVI could accurately detect soybean sudden death syndrome in quadrats of homogeneous soybean canopies with high accuracy (> 0.75) even at early stages of disease development (Raza et al. 2020). However, the permutation importance scores of all image-derived features were very low (< 0.01), and classification accuracy was explained primarily by ancillary data on crop rotation at each sample plot (Raza et al. 2020). We observed similar results for the permutation importance scores of spectral and VI features in all our models. Permutation feature importance scores for all random forest models revealed no consistent pattern in the contribution of specific bands or VIs to classification accuracy. The variation in importance of individual features was high for all classifiers, preventing a stable ranking of potential GDM indicators for either image type (Supplementary Figure S4).

In regions characterized by frequent precipitation, such as the northeastern United States, cloud cover diminishes temporal resolution. *Plasmopara viticola*, the causal agent of GDM, is an aggressive oomycete pathogen with a rapid, polycyclic infection cycle. GDM symptoms can become severe in a matter of days, making high temporal resolution critical for epidemic surveillance. Over the course of this experiment, 3 of 16, 3 of 53, and 5 of 47 PlanetScope images were usable in 2020, 2021, and 2022 respectively (Table 1). The primary cause of the low proportion of usable acquisitions is cloud cover or cloud shadow over the area of interest. This will remain a problem in New York and other cool climate and/or tropical regions where heavy precipitation is common during the grape growing season.

While SkySat offers higher spatial resolution with 0.5-meter pixel size, the capacity for panel-by-panel GDM detection is hindered by the necessity to perform additional co-registration of the imagery and the difficulty of isolating vines from non-targets (soil, weeds, cover crops etc.). Frazier and Hemingway highlight that the positional uncertainty in Planet imagery (up to 10 meters RMSE for the orthorectified image product) necessitates rigorous geometric corrections for applications like precision agriculture and any spatially explicit time-series analysis of vegetation (Frazier and Hemingway 2021). Additionally, both PlanetScope and SkySat satellite constellations are comprised of a fleet of spaceborne sensors, meaning that successive images over the same site are not necessarily acquired by the same sensor on each date. In an independent quality assessment of the SkySat fleet, Saunier et al. report that although signal-to-noise ratios fall within the manufacturer-reported range, the radiometric quality of individual SkySat cameras can differ substantially (Saunier et al. 2022). These sources of variation in geometric and radiometric quality impact the accuracy and reliability of any derived optical data and should be mitigated to the extent possible before imagery is used in decision support systems for crop protection.

Beyond the operational challenges discussed above, disease biology can help explain why GDM detection and surveillance via multispectral satellite imagery is challenging at low incidence and severity levels. Across both data streams assessed, we find that detection capability is closely linked to disease progression. The earliest primary *P. viticola* infections typically occur when rain splash transports oospores that have overwintered in the soil to young grapevine tissue in the lower canopy (Koledenkova et al. 2022; Ash 2000). *Plasmopara viticola* rarely colonizes the upper foliage until several days to weeks later, depending on weather and management intervention (Gessler, Pertot, and Perazzolli 2011). Studies that have successfully used commercial Earth observations for plant disease detection have thus far been limited to diseases affecting the upper canopy (Raza et al. 2020; Guo et al. 2022; Shi et al. 2018). Satellite imagery captures reflectance primarily from leaves in the upper canopy, which do not exhibit chlorotic oil-spot symptoms or defoliation until GDM is further established. The earliest detection of significant differences in any VI in any year between highly diseased and non-diseased vines occurred on 2021-07-26, when median PlanetScope NDVI was lower (P < 0.0001) for pixels corresponding to panels with high incidence and high severity (Figure 4). In 2022, there were no usable SkySat acquisitions after 20-July, but VIs derived from PlanetScope imagery acquired just six days later revealed significant differences between low and high incidence and severity pixels (Figure 3 and Figure 4). This change in detection capability corresponds to increases of 32% in the number of high incidence panels and 43% in high severity panels over the intervening six days (Supplementary Figure S3).

Disease detection in sparse canopies like early season grapevine is further challenged by the contribution of shadow, soil, and other vegetation to the reflectance signal. The substantial soil signal for some grapevine panels could explain why EVI and ARVI, which both incorporate a blue band correction known to mitigate soil background noise, had the strongest correlations with GDM intensity for SkySat images and proved effective at distinguishing between GDM classes (Table 2, Figure 3). It is especially difficult to isolate “pure vine” signal from surrounding vegetation without reference spectra to help distinguish between target and non-target plant tissue. Higher spectral resolution could improve our capacity to isolate vine reflectance from background by taking advantage of finer spectral gradations to unmix multiple endmembers. Recent studies have already demonstrated the utility of spectral unmixing for accurate removal of shadow and soil (Sousa and Small 2018; Galvan et al. 2023). These techniques are increasingly important as more vineyards implement under-vine and inter-row cover crops and as researchers attempt to replicate studies across many field sites with differing cultivation practices. Improved spectral resolution could also enhance VI-based detection at low NDVI steps.

Canopy size impacted our ability to detect differences in VIs between high and low damage classes. Within higher NDVI brackets (NDVI >0.76) we observed higher EVI, GRVI, and ARVI values for healthier panels with respect to diseased panels in 2020 and 2021 (Figure 3, SkySat imagery only). However, this relationship did not hold true at lower NDVI steps. Despite test accuracy and F1 scores both exceeding 0.78, the random forest models trained on three years of PlanetScope and SkySat data performed poorly at classifying high incidence GDM for images acquired during advanced disease stages in 2021 and 2022 (Figure 5). In a comparison of PlanetScope VIs for cover crop yield estimation, Kharel et al. also found that NDVI was positively correlated with biomass across their study period, supporting the link between NDVI and amount of healthy vegetation (Kharel et al. 2023). Further, they observed that correlation coefficients between biomass, VIs, and spectral bands depended on the time of image acquisition, with highest correlation observed at late vegetative growth stages. Like our disease classifiers, Kharel et al.’s random forest models trained on PlanetScope VIs and single bands could not accurately predict biomass even at highly correlated time points. However, the authors highlight that random forest models trained on proximal hyperspectral canopy measurements had higher prediction accuracy (R^2^ of 0.46) (Kharel et al. 2023).

We hypothesized that it is possible to detect GDM infection using multispectral satellite images because the disease elicits symptoms known to cause spectral divergence from healthy plant tissue. It is important to emphasize that GDM infection is one among many abiotic and biotic stressors capable of inducing similar spectral responses, and that increased visible reflectance coupled with lower NIR reflectance are non-specific indicators of vegetation stress. Differentiating between these co-occurring stressors, though unfeasible with multispectral sensing platforms, may be possible with full-spectrum imaging spectroscopy, or hyperspectral imaging, which measures reflectance across a wider range of the electromagnetic spectrum (∼400-2500 nm) at finer spectral intervals (< 10 nm). Recent studies using hyperspectral sensors suggest that asymptomatic and early detection is possible for other grapevine diseases, despite a lack of visible foliar symptoms. Romero Galvan et al. successfully differentiated healthy vines from vines asymptomatically infected with grapevine leafroll virus 3 using AVIRISNG imagery (380-2510 nm spectral range, 5 nm spectral resolution, 1 m spatial resolution) (Galvan et al. 2023). Other studies achieved similar results for asymptomatic leafroll virus detection in the lab or greenhouse (Gao et al. 2020; Bendel et al. 2020; Naidu et al. 2009). Oerke et al. identified asymptomatic/early-stage GDM infections in controlled-environment hyperspectral imaging experiments, indicating the potential for early detection with the addition of reflectance data in the shortwave infrared (SWIR, ∼1200-2500 nm) (Oerke, Herzog, and Toepfer 2016). These wavelengths are sensitive to changes in vegetation water, nutrient, and pigment content, making SWIR imaging a potentially powerful tool for detecting biophysical and biochemical changes that occur when plants undergo biotic and abiotic stress (Asner et al. 2015; Liu et al. 2021; Sobejano-Paz et al. 2020; Sapes et al. 2021; Serbin et al. 2014). Although *P. viticola* alters pigment concentrations, sugar translocation, photosynthesis, and water uptake immediately upon infection, the limited spectral range and resolution of SkySat and PlanetScope imagery is insufficient to detect these subtle, non-visible changes.

Of the two metrics used here to assess GDM damage, incidence and severity, incidence may provide a more consistent measure of disease intensity. GDM is characterized by visually distinct signs and symptoms (Figure 1), such that counting visibly infected units (i.e. incidence rating) may be more straightforward and yield more consistent results than attempting to precisely quantify the symptomatic area of an infected unit (i.e. severity rating). Indeed, we found that VIs differ significantly (P < 0.05) more often between GDM incidence classes than between severity classes (Figures 3 and 4). Random forest incidence classifiers also had higher accuracy scores than severity classifiers across image datasets (Tables 4 and 5). We would be remiss not to acknowledge the role scouting bias could play in the variable capacity observed across years in our study. It is well established that inter- and intra-rater variability can have an outsized impact on the accuracy and precision of plant disease incidence and severity estimation (Nutter and Schultz 1995; Bock et al. 2010). Although standard area diagrams and rigorous scout training have been shown to improve reliability of severity ratings, inherent rater subjectivity and the influence of environment, among other factors, remain a challenge (Bock et al. 2022; Del Ponte et al. 2017). In our study, some members of the disease rating team differed across years and, though training and measurement protocols were standardized, ratings cannot be guaranteed to be free from bias. Additionally, the weekly random sample of twenty leaves is a technical, rather than biological, replicate, such that GDM is quantified on a different subset of leaves each week.

Leveraging proximal imaging spectrometers to complement multispectral satellite platforms could help relate fine-scale, disease-induced changes in canopy reflectance to the relatively coarser signal observed by satellites, as well as mitigate error due to human bias. To address this challenge, Liu et al. built and validated a computer-vision based GDM detection pipeline that uses geo-referenced, side-canopy images acquired by an autonomous rover to quantify symptom severity (Liu et al. 2022). By standardizing and automating severity estimation, such systems can reduce potential error caused by human rater bias. Future work must evaluate whether and how much autonomous GDM ratings improve accuracy and precision relative to the human standard, and to what extent the rover validation data increases agreement between satellite-derived and proximal estimates of disease severity. Future work with paired proximal and remote sensing could explore adding weather conditions and relevant management actions to test whether these features improve classification accuracy for GDM incidence and severity.

The goal of our work was to determine if we can detect and track relative GDM damage in a research vineyard with an artificial disease gradient established via variable fungicide application. Our results indicate that high resolution, multispectral satellite imagery coupled with supervised machine learning methods, though undeniably promising, are not yet able to produce maps of GDM damage sufficiently reliable to inform rapid management intervention. Both satellite data streams we assessed have potential to contribute to grapevine health surveillance, but users should be aware of the possibilities and limitations associated with each. We found that late-stage GDM detection and season-long surveillance are hindered by insufficient temporal coverage due to cloud cover, low spectral resolution of currently available satellite image products, and the confounding influence of reflectance from soil, shadow, and non-target vegetation. Additionally, while we specifically sought to control other pathogens in the vineyard, other diseases did occur at low levels. Improved spectral unmixing algorithms and forthcoming hyperspectral satellite platforms, such as Planet Lab’s Pelican and Carbonmapper systems (“NASA-Built Greenhouse Gas Detector Moves Closer to Launch” 2023; “Planet Pelican” 2022) and Pixxel’s imaging suite (“Pixxel Hyperspectral Imaging Constellation” 2023) will hopefully help resolve these challenges, as well as enable differentiating GDM from co-occurring abiotic stressors or from other grapevine diseases.

## V. Acknowledgements

This research was funded by the Cornell Institute for Digital Agriculture Research Venture Fund and USDA NIFA Small Farms Program (2022-68006-36148). Many thanks to Holly Staid, Perry Wivell, Megan Walker, Monique Michaud, Jonas Campagna, Eric Winarski, Fernando Romero Galvan, Bruno Aragon, Rocío Calderón, Manushi Trivedi, and Ying Sun for their support in data collection and analysis.

## Supplementary Tables

**Table S1.** Mean and maximum GDM severity and incidence, 2020-2022.

| GDM Severity |  |  |
| --- | --- | --- |
| Year | Mean end-of-season (%) | Maximum (%) |
| 2020 | 3 | 82 |
| 2021 | 6 | 81 |
| 2022 | 14 | 57 |
| GDM Incidence |  |  |
| Year | Mean end-of-season (%) | Maximum (%) |
| 2020 | 45 | 100 |
| 2021 | 22 | 100 |
| 2022 | 80 | 100 |

## Literature cited

Aragon, Bruno, Matteo G. Ziliani, Rasmus Houborg, Trenton E. Franz, and Matthew F. McCabe. 2021. “CubeSats Deliver New Insights into Agricultural Water Use at Daily and 3 m Resolutions.” Scientific Reports 11 (1): 12131. 10.1038/s41598-021-91646-w.

Ash, G. 2000. “Downy Mildew of Grape.” The Plant Health Instructor. 10.1094/phi-i-2000-1112-01.

Asner, Gregory P., Roberta E. Martin, Christopher B. Anderson, and David E. Knapp. 2015. “Quantifying Forest Canopy Traits: Imaging Spectroscopy versus Field Survey.” Remote Sensing of Environment 158: 15–27. 10.1016/j.rse.2014.11.011.

Baloloy, Alvin Balidoy, Ariel Conferido Blanco, Christian Gumbao Candido, Reginal Jay Labadisos Argamosa, John Bart Lovern Caboboy Dumalag, Lady Lee Carandang Dimapilis, and Enrico Camero Paringit. 2018. “Estimation of Mangrove Forest Aboveground Biomass Using Multispectral Bands, Vegetation Indices and Biophysical Variables Derived From Optical Satellite Imageries: Rapideye, PlanetScope and Sentinel-2.” ISPRS Annals of the Photogrammetry, Remote Sensing and Spatial Information Sciences IV–3 (3): 29–36. 10.5194/isprs-annals-IV-3-29-2018.

Barreto, Abel, Facundo Ramón Ispizua Yamati, Mark Varrelmann, Stefan Paulus, and Anne Katrin Mahlein. 2023. “Disease Incidence and Severity of Cercospora Leaf Spot in Sugar Beet Assessed by Multispectral Unmanned Aerial Images and Machine Learning.” Plant Disease 107 (1): 188–200. 10.1094/PDIS-12-21-2734-RE.

Bendel, Nele, Anna Kicherer, Andreas Backhaus, Janine Köckerling, Michael Maixner, Elvira Bleser, Hans-Christian Klück, Udo Seiffert, Ralf T. Voegele, and Reinhard Töpfer. 2020. “Detection of Grapevine Leafroll-Associated Virus 1 and 3 in White and Red Grapevine Cultivars Using Hyperspectral Imaging.” Remote Sensing 12 (10): 1693. 10.3390/rs12101693.

Bock, C. H., G. H. Poole, P.E. Parker, and T. Gottwald. 2010. “Critical Reviews in Plant Sciences Plant Disease Severity Estimated Visually, by Digital Photography and Image Analysis, and by Hyperspectral Imaging Plant Disease Severity Estimated Visually, by Digital Photography and Image Analysis, and by Hyperspectral Imaging.” Critical Reviews in Plant Sciences 29: 59–107. 10.1080/07352681003617285.

Bock, Clive H., Jayme G. A. Barbedo, Anne-Katrin Mahlein, and Emerson M. Del Ponte. 2022. “A Special Issue on Phytopathometry — Visual Assessment, Remote Sensing, and Artificial Intelligence in the Twenty-First Century.” Tropical Plant Pathology 47 (1): 1–4. 10.1007/s40858-022-00498-w.

Calderón, Rocío, M. Montes-Borrego, B. B. Landa, J. A. Navas-Cortés, and P. J. Zarco-Tejada. 2014. “Detection of Downy Mildew of Opium Poppy Using High-Resolution Multi-Spectral and Thermal Imagery Acquired with an Unmanned Aerial Vehicle.” Precision Agriculture 15 (6): 639–61. 10.1007/s11119-014-9360-y.

Calderón, Rocío, Juan A. Navas-Cortés, and Pablo J. Zarco-Tejada. 2015. “Early Detection and Quantification of Verticillium Wilt in Olive Using Hyperspectral and Thermal Imagery over Large Areas.” Remote Sensing 7 (5): 5584–5610. 10.3390/rs70505584.

Camargo, Meyriele P., Bruna V. Momesso, Marlon H. Hahn, and Henrique S. S. Duarte. 2019. “Development and Validation of a Standard Area Diagram Set to Estimate Severity of Grapevine Downy Mildew on Vitis Labrusca.” European Journal of Plant Pathology 155 (3): 1033–38. 10.1007/s10658-019-01806-y.

Campbell, Sarah E., Phillip M. Brannen, Harald Scherm, Nathan Eason, and Clark MacAllister. 2021. “Efficacy of Fungicide Treatments for Plasmopara Viticola Control and Occurrence of Strobilurin Field Resistance in Vineyards in Georgia, USA.” Crop Protection 139 (January). 10.1016/j.cropro.2020.105371.

Carlson, Gerald A., and J.C. Headley. 1987. “Economic Aspects of Integrated Pest Management Threshold Determination.” Plant Disease 71: 459–62.

Corio-Costet, M. F. 2012. “Fungicide Resistance in *Plasmopara Viticola* in France and Anti-Resistance Measures.” In Fungicide Resistance in Crop Protection: Risk and Management, 157–71. UK: CABI. 10.1079/9781845939052.0157.

Cséfalvay, Ladislav, Gabriele Di Gaspero, Karel Matouš, Diana Bellin, Benedetto Ruperti, and Julie Olejníčková. 2009. “Pre-Symptomatic Detection of Plasmopara Viticola Infection in Grapevine Leaves Using Chlorophyll Fluorescence Imaging.” European Journal of Plant Pathology 125 (2): 291–302. 10.1007/s10658-009-9482-7.

Curran, Paul J. 1989. “Remote Sensing of Foliar Chemistry.” Remote Sensing of Environment 30 (3): 271–78. 10.1016/0034-4257(89)90069-2.

Daniel, Wayne W. 1990. Applied Nonparametric Statistics. 2nd ed. Boston: PWS-KENT Pub.

Du, Jinyang, John S. Kimball, Rajat Bindlish, Jeffrey P. Walker, and Jennifer D. Watts. 2022. “Local Scale (3-m) Soil Moisture Mapping Using SMAP and Planet SuperDove.” Remote Sensing 14 (15): 3812. 10.3390/rs14153812.

“Duro Product Summary.” 2021. https://www.swiftnav.com/sites/default/files/duro_product_summary.pdf.

Feng, Xuewen, and Anton Baudoin. 2018. “First Report of Carboxylic Acid Amide Fungicide Resistance in *Plasmopara Viticola* (Grapevine Downy Mildew) in North America.” Plant Health Progress 19 (2): 139–139. 10.1094/PHP-01-18-0005-BR.

Frazier, Amy E., and Benjamin L. Hemingway. 2021. “A Technical Review of Planet Smallsat Data: Practical Considerations for Processing and Using PlanetScope Imagery.” Remote Sensing 2021, Vol. 13, Page 3930 13 (19): 3930. 10.3390/RS13193930.

Galvan, Fernando E. Romero, Ryan Pavlick, Graham Trolley, Somil Aggarwal, Daniel Sousa, Charles Starr, Elisabeth Forrestel, et al. 2023. “Scalable Early Detection of Grapevine Viral Infection with Airborne Imaging Spectroscopy.” Phytopathology 113 (8): 1439–46. 10.1094/PHYTO-01-23-0030-R.

Gamon, J.A., J. Peñuelas, and C.B. Field. 1992. “A Narrow-Waveband Spectral Index That Tracks Diurnal Changes in Photosynthetic Efficiency.” Remote Sensing of Environment 41 (1): 35–44. 10.1016/0034-4257(92)90059-S.

Gao, Zongmei, Lav R. Khot, Rayapati A. Naidu, and Qin Zhang. 2020. “Early Detection of Grapevine Leafroll Disease in a Red-Berried Wine Grape Cultivar Using Hyperspectral Imaging.” Computers and Electronics in Agriculture 179 (December): 105807. 10.1016/j.compag.2020.105807.

Gennaro, Salvatore F. di, Enrico Battiston, Stefano di Marco, Osvaldo Facini, Alessandro Matese, Marco Nocentini, Alberto Palliotti, and Laura Mugnai. 2016. “Unmanned Aerial Vehicle (UAV)-Based Remote Sensing to Monitor Grapevine Leaf Stripe Disease within a Vineyard Affected by Esca Complex.” Phytopathologia Mediterranea 55 (2): 262–75. 10.14601/Phytopathol_Mediterr-18312.

Gessler, Cesare, Ilaria Pertot, and Michele Perazzolli. 2011. “Plasmopara Viticola: A Review of Knowledge on Downy Mildew of Grapevine and Effective Disease Management.” Phytopathologia Mediterranea 50 (1): 3–44. 10.14601/Phytopathol_Mediterr-9360.

Giovos, Rigas, Dimitrios Tassopoulos, Dionissios Kalivas, Nestor Lougkos, and Anastasia Priovolou. 2021. “Remote Sensing Vegetation Indices in Viticulture: A Critical Review.” Agriculture (Switzerland). MDPI AG. 10.3390/agriculture11050457.

Gisi, Ulrich, and Helge Sierotzki. 2008. “Fungicide Modes of Action and Resistance in Downy Mildews.” In European Journal of Plant Pathology, 122:157–67. 10.1007/s10658-008-9290-5.

Glenn, Edward, Alfredo Huete, Pamela Nagler, and Stephen Nelson. 2008. “Relationship Between Remotely-Sensed Vegetation Indices, Canopy Attributes and Plant Physiological Processes: What Vegetation Indices Can and Cannot Tell Us About the Landscape.” Sensors 8 (4): 2136–60. 10.3390/s8042136.

Gold, Kaitlin M., Philip A. Townsend, Ittai Herrmann, and Amanda J. Gevens. 2020. “Investigating Potato Late Blight Physiological Differences across Potato Cultivars with Spectroscopy and Machine Learning.” Plant Science 295 (October 2019): 110316. 10.1016/j.plantsci.2019.110316.

Guo, Jiawei, Yu Jin, Huichun Ye, Wenjiang Huang, Jinling Zhao, Bei Cui, Fucheng Liu, and Jiajian Deng. 2022. “Recognition of Areca Leaf Yellow Disease Based on Planetscope Satellite Imagery.” Agronomy 12 (1). 10.3390/agronomy12010014.

Haboudane, Driss, John R. Miller, Nicolas Tremblay, Pablo J. Zarco-Tejada, and Louise Dextraze. 2002. “Integrated Narrow-Band Vegetation Indices for Prediction of Crop Chlorophyll Content for Application to Precision Agriculture.” Remote Sensing of Environment 81 (2–3): 416–26. 10.1016/S0034-4257(02)00018-4.

Hassanzadeh, Amirhossein, Fei Zhang, Sean Murphy, Sarah Pethybridge, and Jan Van Aardt. 2021. “Toward Crop Maturity Assessment via UAS-Based Imaging Spectroscopy - A Snap Bean Pod Size Classification Field Study.” IEEE Transactions on Geoscience and Remote Sensing 14 (8). 10.1109/TGRS.2021.3134564.

Heim, René, Ian Wright, Peter Scarth, Angus Carnegie, Dominique Taylor, and Jens Oldeland. 2019. “Multispectral, Aerial Disease Detection for Myrtle Rust (Austropuccinia Psidii) on a Lemon Myrtle Plantation.” Drones 3 (1): 25. 10.3390/drones3010025.

Helman, David, Idan Bahat, Yishai Netzer, Alon Ben-Gal, Victor Alchanatis, Aviva Peeters, and Yafit Cohen. 2018. “Using Time Series of High-Resolution Planet Satellite Images to Monitor Grapevine Stem Water Potential in Commercial Vineyards.” Remote Sensing 10 (10): 1615. 10.3390/rs10101615.

Huang, Sha, Lina Tang, Joseph P. Hupy, Yang Wang, and Guofan Shao. 2021. “A Commentary Review on the Use of Normalized Difference Vegetation Index (NDVI) in the Era of Popular Remote Sensing.” Journal of Forestry Research. Northeast Forestry University. 10.1007/s11676-020-01155-1.

Huete, A., C. Justice, and H. Liu. 1994. “Development of Vegetation and Soil Indices for MODIS-EOS.” Vol. 49.

Iordache, Marian Daniel, Vasco Mantas, Elsa Baltazar, Klaas Pauly, and Nicolas Lewyckyj. 2020. “A Machine Learning Approach to Detecting Pine Wilt Disease Using Airborne Spectral Imagery.” Remote Sensing 12 (14). 10.3390/RS12142280.

Jackson, R. D. 1986. “Remote Sensing of Biotic and Abiotic Plant Stress.” Annual Review of Phytopathology 24 (1): 265–87. 10.1146/annurev.py.24.090186.001405.

Kaufman, Y.J., and D. Tanre. 1992. “Atmospherically Resistant Vegetation Index (ARVI) for EOS-MODIS.” IEEE Transactions on Geoscience and Remote Sensing 30 (2): 261–70. 10.1109/36.134076.

Khanal, Sami, Kushal KC, John P. Fulton, Scott Shearer, and Erdal Ozkan. 2020. “Remote Sensing in Agriculture—Accomplishments, Limitations, and Opportunities.” Remote Sensing 12 (22): 3783. 10.3390/rs12223783.

Kharel, Tulsi P., Ammar B. Bhandari, Partson Mubvumba, Heather L. Tyler, Reginald S. Fletcher, and Krishna N. Reddy. 2023. “Mixed-Species Cover Crop Biomass Estimation Using Planet Imagery.” Sensors 23 (3): 1541. 10.3390/s23031541.

Kim, M., S. Park, C. Anderson, and G.L. Stensaas. 2022. “System Characterization Report on Planet’s SuperDove.”

Knipling, Edward B. 1970. “Physical and Physiological Basis for the Reflectance of Visible and Near-Infrared Radiation from Vegetation.” Remote Sensing of Environment 1 (3): 155–59. 10.1016/S0034-4257(70)80021-9.

Koledenkova, Kseniia, Qassim Esmaeel, Cédric Jacquard, Jerzy Nowak, Christophe Clément, and Essaid Ait Barka. 2022. “Plasmopara Viticola the Causal Agent of Downy Mildew of Grapevine: From Its Taxonomy to Disease Management.” Frontiers in Microbiology. Frontiers Media S.A. 10.3389/fmicb.2022.889472.

Lacotte, Virginie, Sergio Peignier, Marc Raynal, Isabelle Demeaux, François Delmotte, and Pedro da Silva. 2022. “Spatial–Spectral Analysis of Hyperspectral Images Reveals Early Detection of Downy Mildew on Grapevine Leaves.” International Journal of Molecular Sciences 23 (17). 10.3390/ijms231710012.

Lemaitre, Guillaume, Fernando Nogueira, and Aridas K. Christos. 2017. “Imbalanced-Learn: A Python Toolbox to Tackle the Curse of Imbalanced Datasets in Machine Learning.” Journal of Machine Learning Research 18: 1–5.

Liu, Ertai, Kaitlin M. Gold, David Combs, Lance Cadle-Davidson, and Yu Jiang. 2022. “Deep Semantic Segmentation for the Quantification of Grape Foliar Diseases in the Vineyard.” Frontiers in Plant Science 13 (September). 10.3389/fpls.2022.978761.

Liu, Hui Qing, and Alfred0 Huete. 1995. “A Feedback Based Modification of the NDVI to Minimize Canopy Background and Atmospheric Noise.” IEEE TRANSACTIONS ON GEOSCIENCE AND REMOTE SENSING. Vol. 33.

Liu, Nanfeng, Philip A. Townsend, Mack R. Naber, Paul C. Bethke, William B. Hills, and Yi Wang. 2021. “Hyperspectral Imagery to Monitor Crop Nutrient Status within and across Growing Seasons.” Remote Sensing of Environment 255 (August 2020): 112303. 10.1016/j.rse.2021.112303.

MacDonald, Sarah L., Matthew Staid, Melissa Staid, and Monica L. Cooper. 2016. “Remote Hyperspectral Imaging of Grapevine Leafroll-Associated Virus 3 in Cabernet Sauvignon Vineyards.” Computers and Electronics in Agriculture 130 (November): 109–17. 10.1016/j.compag.2016.10.003.

Mann, H. B., and D R Whitney. 1947. “On a Test of Whether One of Two Random Variables Is Stochastically Larger than the Other.” The Annals of Mathematical Statistics 18 (1): 50 – 60. 10.1214/aoms/1177730491.

Matese, Alessandro, Salvatore Filippo Di Gennaro, Giorgia Orlandi, Matteo Gatti, and Stefano Poni. 2022. “Assessing Grapevine Biophysical Parameters From Unmanned Aerial Vehicles Hyperspectral Imagery.” Frontiers in Plant Science 13 (June). 10.3389/fpls.2022.898722.

Meyers, James M., Nick Dokoozlian, Casey Ryan, Cella Bioni, and Justine E. Vanden Heuvel. 2020. “A New, Satellite NDVI-Based Sampling Protocol for Grape Maturation Monitoring.” Remote Sensing 12 (7). 10.3390/rs12071159.

Moran, M.S., Y. Inoue, and E.M. Barnes. 1997. “Opportunities and Limitations for Image-Based Remote Sensing in Precision Crop Management.” Remote Sensing of Environment 61 (3): 319–46. 10.1016/S0034-4257(97)00045-X.

Nagarajan, S., G. Seibold, J. Kranz, E. E. Saari, and L. M. Joshi. 1984. “Monitoring Wheat Rust Epidemics with the Landsat-2 Satellite [Triticum Aestivum and Triticum Durum Infected with Puccinia Striiformis and Puccinia Recondita f. Sp. Tritici].” Phytopathology.

Naidu, Rayapati A., Eileen M. Perry, Francis J. Pierce, and Tefera Mekuria. 2009. “The Potential of Spectral Reflectance Technique for the Detection of Grapevine Leafroll-Associated Virus-3 in Two Red-Berried Wine Grape Cultivars.” Computers and Electronics in Agriculture 66 (1): 38–45. 10.1016/j.compag.2008.11.007.

“NASA-Built Greenhouse Gas Detector Moves Closer to Launch.” 2023. NASA Jet Propulsion Laboratory. September 14, 2023. https://www.jpl.nasa.gov/news/nasa-built-greenhouse-gas-detector-moves-closer-to-launch.

Nilsson, H. 1995. “Remote Sensing and Image Analysis in Plant Pathology.” Annual Review of Phytopathology 33 (1): 489–528. 10.1146/annurev.py.33.090195.002421.

Nutter, F. W., Tylka, G. L., Guan, J., Moreira, A. J., Marett, C. C., Rosburg, T. R., Basart, J. P., & Chong, C. S. (2002). “Use of remote sensing to detect soybean cyst nematode-induced plant stress.” Journal of Nematology 34(3), 222–231.

Nutter, Forrest W., and Patricia M. Schultz. 1995. “Improving the Accuracy and Precision of Disease Assessments: Selection of Methods and Use of Computer-Aided Training Programs.” Canadian Journal of Plant Pathology 17 (2): 174–84. 10.1080/07060669509500709.

Oerke, Erich Christian, Katja Herzog, and Reinhard Toepfer. 2016. “Hyperspectral Phenotyping of the Reaction of Grapevine Genotypes to Plasmopara Viticola.” Journal of Experimental Botany 67 (18): 5529–43. 10.1093/jxb/erw318.

Pedregosa, F., G. Varoquaux, A. Gramfort, V. Michel, B. Thirion, O. Grisel, M. Blondel, et al. 2011. “Scikit-Learn: Machine Learning in Python.” Journal of Machine Learning Research 12: 2825–30.

“Pixxel Hyperspectral Imaging Constellation.” 2023. EO Portal. June 7, 2023. https://www.eoportal.org/satellite-missions/pixxel#sensorcomplement.

“Planet Imagery Product Specifications.” 2022. https://assets.planet.com/docs/Planet_Combined_Imagery_Product_Specs_letter_screen.pdf?_gl=1*m631iy*_gcl_aw*R0NMLjE2OTQxMTc1MDQuQ2p3S0NBanc2ZVduQmhBS0Vpd0FEcG53OW1oZ0Vyb0xRZTQtQTZzOTE4NlBfSGhySUo1VjJTcHFqOUozVzVLdkNYbUdkRnhxcHdLWnpSb0NjYmdRQXZEX0J3RQ..*_gcl_au*MTIwMzY0MDc4NS4xNjk0MTE3NTA0.

“Planet Pelican.” 2022. EO Portal. May 12, 2022. https://www.eoportal.org/satellite-missions/planet-pelican#eop-quick-facts-section.

Ponte, Emerson M. Del, Sarah J. Pethybridge, Clive H. Bock, Sami J. Michereff, Franklin J. Machado, and Piérri Spolti. 2017. “Standard Area Diagrams for Aiding Severity Estimation: Scientometrics, Pathosystems, and Methodological Trends in the Last 25 Years.” Phytopathology 107 (10): 1161–74. 10.1094/PHYTO-02-17-0069-FI.

Qi, J., A. Chehbouni, A.R. Huete, Y.H. Kerr, and S. Sorooshian. 1994. “A Modified Soil Adjusted Vegetation Index.” Remote Sensing of Environment 48 (2): 119–26. 10.1016/0034-4257(94)90134-1.

Raza, Muhammad M., Chris Harding, Matt Liebman, and Leonor F. Leandro. 2020. “Exploring the Potential of High-Resolution Satellite Imagery for the Detection of Soybean Sudden Death Syndrome.” Remote Sensing 12 (7). 10.3390/rs12071213.

Rouse Jr., J.W., R.H. Haas, J.A. Schell, and D.W. Deering. 1974. “Monitoring Vegetation Systems in the Great Plains with ERTS.” In NASA Goddard Space Flight Center 3d ERTS-1 Symp. https://ntrs.nasa.gov/citations/19740022614.

Santos, Ricardo F., Bart A. Fraaije, Lucas da R. Garrido, Cláudia B. Monteiro-Vitorello, and Lilian Amorim. 2020. “Multiple Resistance of *Plasmopara Viticola* to QoI and CAA Fungicides in Brazil.” Plant Pathology 69 (9): 1708–20. 10.1111/ppa.13254.

Sapes, Gerard, Cathleen Lapadat, Anna K Scweiger, Jennifer Juzwik, Rebecca Montgomery, Hamed Gholizadeh, Philip Townsend, John Gamon, and Jeannine Cavender-Bares. 2021. “Canopy Spectral Reflectance Detects Oak Wilt at the Landscape Scale Using Phylogenetic Discrimination.” BioRxiv, no. February: 1–49. 10.1016/j.rse.2022.112961.

Saunier, Sebastien, Gizem Karakas, Ilyas Yalcin, Fay Done, Rubinder Mannan, Clement Albinet, Philippe Goryl, and Sultan Kocaman. 2022. “SkySat Data Quality Assessment within the EDAP Framework.” Remote Sensing 14 (7). 10.3390/rs14071646.

Sawyer, Erica, Eve Laroche-Pinel, Madison Flasco, Monica L. Cooper, Benjamin Corrales, Marc Fuchs, and Luca Brillante. 2023. “Phenotyping Grapevine Red Blotch Virus and Grapevine Leafroll-Associated Viruses before and after Symptom Expression through Machine-Learning Analysis of Hyperspectral Images.” Frontiers in Plant Science 14 (March). 10.3389/fpls.2023.1117869.

Scheffler, Daniel, André Hollstein, Hannes Diedrich, Karl Segl, and Patrick Hostert. 2017. “AROSICS: An Automated and Robust Open-Source Image Co-Registration Software for Multi-Sensor Satellite Data.” Remote Sensing 9 (7). 10.3390/rs9070676.

Serbin, Shawn P., Aditya Singh, Brenden E. McNeil, Clayton C. Kingdon, and Philip A. Townsend. 2014. “Spectroscopic Determination of Leaf Morphological and Biochemical Traits for Northern Temperate and Boreal Tree Species.” Ecological Applications 24 (7): 1651–69. 10.1890/13-2110.1.

Serbin, Shawn P., and Philip A. Townsend. 2020. “Scaling Functional Traits from Leaves to Canopies.” In Remote Sensing of Plant Biodiversity, edited by Jeannine Cavender-Bares, John A. Gamon, and Philip A. Townsend, 43–82. Springer International Publishing. 10.1007/978-3-030-33157-3.

Sharma, N., L. Heger, D. Combs, K. Gold, and T. D. Miles. 2022. “Assessment of QoI and CAA Fungicide Resistance of *Plasmopara Viticola* Populations in Vineyards of the Great Lakes Region in the United States of America.” BIO Web of Conferences 50 (August): 02011. 10.1051/bioconf/20225002011.

Shi, Yue, Wenjiang Huang, Huichun Ye, Chao Ruan, Naichen Xing, Yun Geng, Yingying Dong, and Dailiang Peng. 2018. “Partial Least Square Discriminant Analysis Based on Normalized Two-Stage Vegetation Indices for Mapping Damage from Rice Diseases Using PlanetScope Datasets.” 10.3390/s18061901.

Sims, Daniel A, and John A Gamon. 2002. “Relationships between Leaf Pigment Content and Spectral Reflectance across a Wide Range of Species, Leaf Structures and Developmental Stages.” Remote Sensing of Environment 81 (2–3): 337–54. 10.1016/S0034-4257(02)00010-X.

Sobejano-Paz, Verónica, Teis Nørgaard Mikkelsen, Andreas Baum, Xingguo Mo, Suxia Liu, Christian Josef Köppl, Mark S. Johnson, Lorant Gulyas, and Mónica García. 2020. “Hyperspectral and Thermal Sensing of Stomatal Conductance, Transpiration, and Photosynthesis for Soybean and Maize under Drought.” Remote Sensing 12 (19): 1–32. 10.3390/RS12193182.

Sousa, Daniel, and Christopher Small. 2018. “Multisensor Analysis of Spectral Dimensionality and Soil Diversity in the Great Central Valley of California.” Sensors (Switzerland) 18 (2). 10.3390/s18020583.

Szabó, Loránd, Dávid Abriha, Kwanele Phinzi, and Szilárd Szabó. 2021. “Urban Vegetation Classification with High-Resolution PlanetScope and SkySat Multispectral Imagery.” Landscape & Environment 15 (1): 66–75. 10.21120/le/15/1/9.

Toffolatti, Silvia L, Giuseppe Russo, Paola Campia, Piero A Bianco, Paolo Borsa, Mauro Coatti, Stefano FF Torriani, and Helge Sierotzki. 2018. “A Time-course Investigation of Resistance to the Carboxylic Acid Amide Mandipropamid in Field Populations of *Plasmopara Viticola* Treated with Anti-resistance Strategies.” Pest Management Science 74 (12): 2822–34. 10.1002/ps.5072.

Tu, Yu-Hsuan, Kasper Johansen, Bruno Aragon, Marcel M. El Hajj, and Matthew F. McCabe. 2022. “The Radiometric Accuracy of the 8-Band Multi-Spectral Surface Reflectance from the Planet SuperDove Constellation.” International Journal of Applied Earth Observation and Geoinformation 114 (November): 103035. 10.1016/j.jag.2022.103035.

Tucker, Compton J. 1979. “Red and Photographic Infrared Linear Combinations for Monitoring Vegetation.” Remote Sensing of Environment 8 (2): 127–50. 10.1016/0034-4257(79)90013-0.

Vitrack-Tamam, Snir, Lilach Holtzman, Reut Dagan, Shai Levi, Yuval Tadmor, Tamir Azizi, Onn Rabinovitz, Amos Naor, and Oded Liran. 2020. “Random Forest Algorithm Improves Detection of Physiological Activity Embedded within Reflectance Spectra Using Stomatal Conductance as a Test Case.” Remote Sensing 12 (14). 10.3390/rs12142213.

Wang, Jing, Dedi Yang, Matteo Detto, Bruce W. Nelson, Min Chen, Kaiyu Guan, Shengbiao Wu, Zhengbing Yan, and Jin Wu. 2020. “Multi-Scale Integration of Satellite Remote Sensing Improves Characterization of Dry-Season Green-up in an Amazon Tropical Evergreen Forest.” Remote Sensing of Environment 246 (September): 111865. 10.1016/J.RSE.2020.111865.

Wang, Junming, Theodore W. Sammis, Vincent P. Gutschick, Mekonnen Gebremichael, Sam O. Dennis, and Robert E. Harrison. 2010. “Review of Satellite Remote Sensing Use in Forest Health Studies∼!2010-01-27∼!2010-04-05∼!2010-06-29∼!” The Open Geography Journal 3 (1): 28–42. 10.2174/1874923201003010028.

Wang, Zhihui, Tiejun Wang, Roshanak Darvishzadeh, Andrew K. Skidmore, Simon Jones, Lola Suarez, William Woodgate, Uta Heiden, Marco Heurich, and John Hearne. 2016. “Vegetation Indices for Mapping Canopy Foliar Nitrogen in a Mixed Temperate Forest.” Remote Sensing 2016, Vol. 8, Page 491 8 (6): 491. 10.3390/RS8060491.

Wilcox, Wayne F., Walter D. Gubler, and Jerry K. Uyemoto, eds. 2015. Compendium of Grape Diseases Disorders and Pests. 2nd ed. St. Paul Minnesota: APS Press.

Yang, Chenghai. 2018. “High Resolution Satellite Imaging Sensors for Precision Agriculture.” Frontiers of Agricultural Science and Engineering. Higher Education Press Limited Company. 10.15302/J-FASE-2018226.

Yao, Haibo, Yanbo Huang, Zuzana Hruska, Steven J. Thomson, and Krishna N. Reddy. 2012. “Using Vegetation Index and Modified Derivative for Early Detection of Soybean Plant Injury from Glyphosate.” Computers and Electronics in Agriculture 89 (November): 145–57. 10.1016/j.compag.2012.09.001.

Yin, Gaofei, Aleixandre Verger, Adrià Descals, Iolanda Filella, and Josep Peñuelas. 2022. “A Broadband Green-Red Vegetation Index for Monitoring Gross Primary Production Phenology.” Journal of Remote Sensing 2022 (January). 10.34133/2022/9764982.

Zeng, Yelu, Dalei Hao, Alfredo Huete, Benjamin Dechant, Joe Berry, Jing M. Chen, Joanna Joiner, et al. 2022. “Optical Vegetation Indices for Monitoring Terrestrial Ecosystems Globally.” Nature Reviews Earth & Environment 3 (7): 477–93. 10.1038/s43017-022-00298-5.

