## Supplementary figures and images for "Assessing the capacity of high-resolution commercial satellite imagery for grapevine downy mildew detection and surveillance in New York state"

### S1

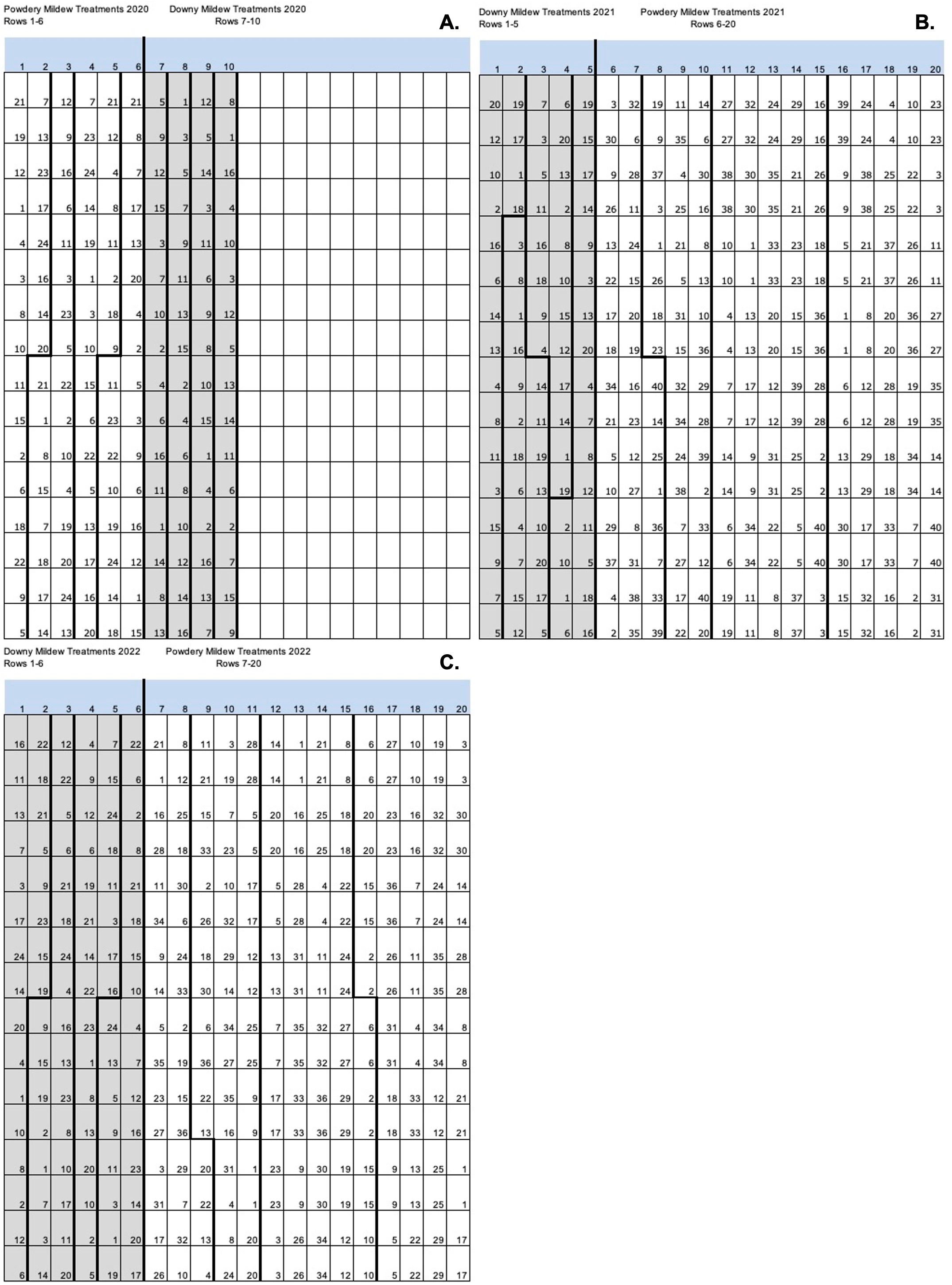

### S2

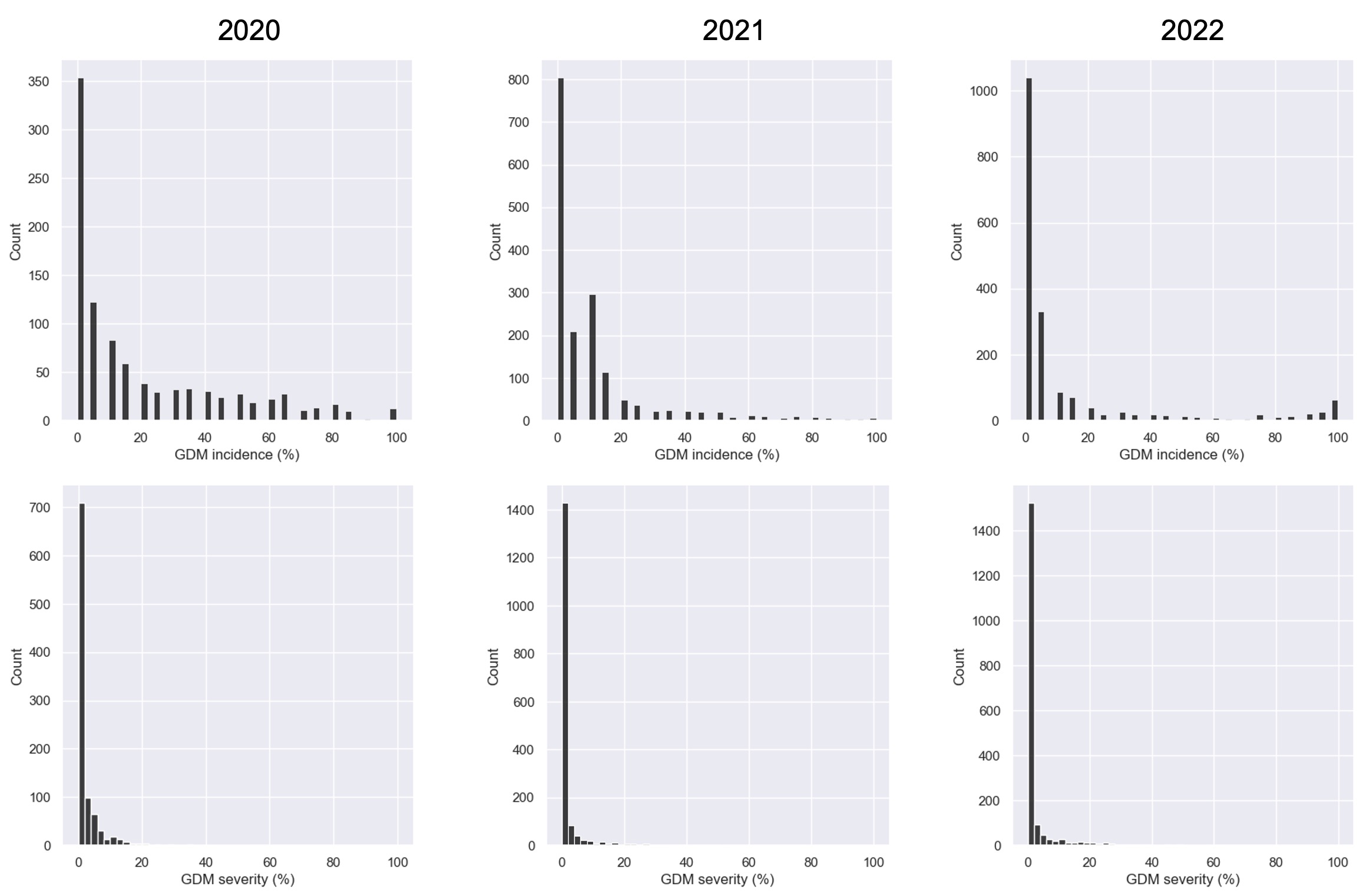

### S3

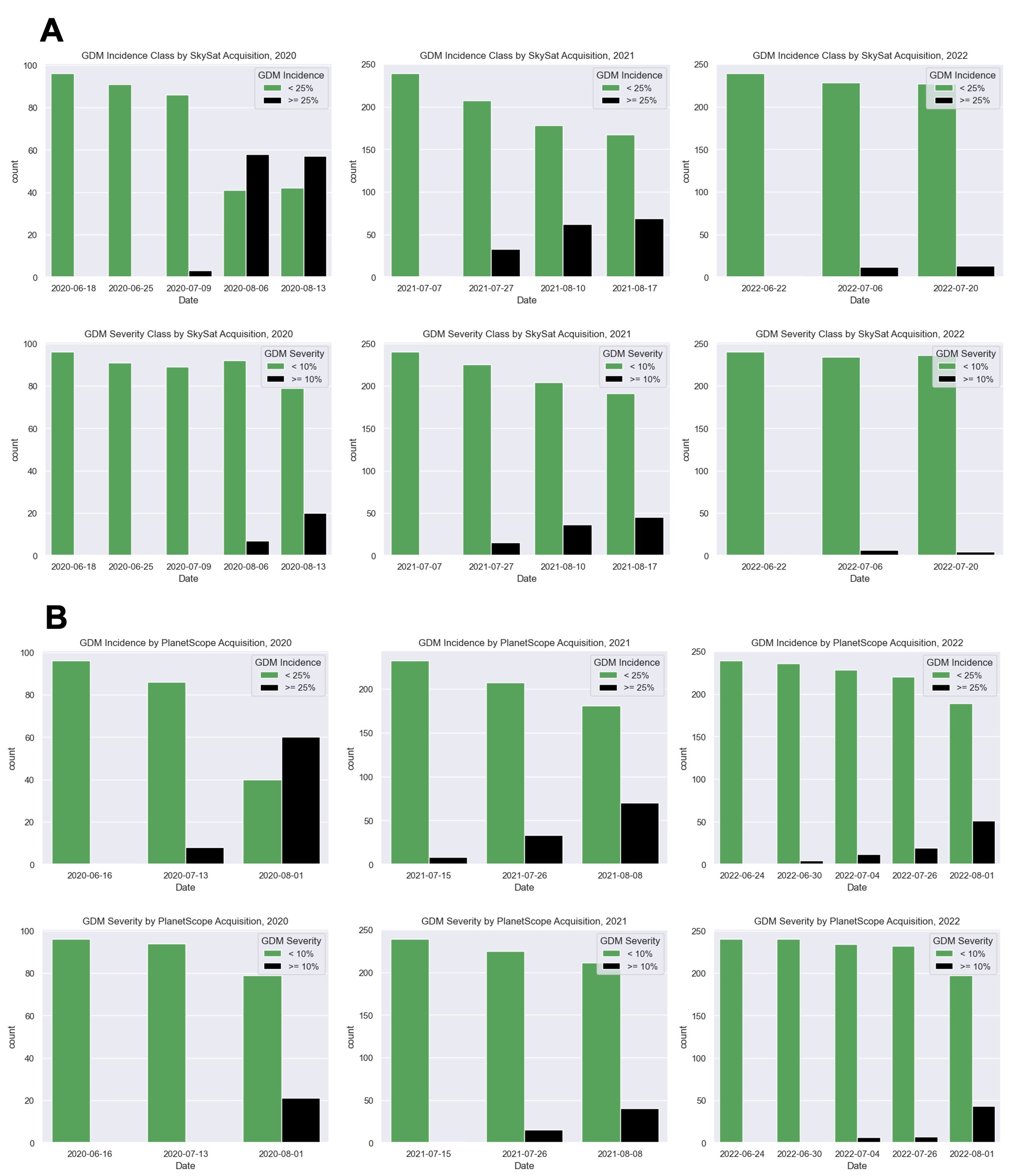

### S4

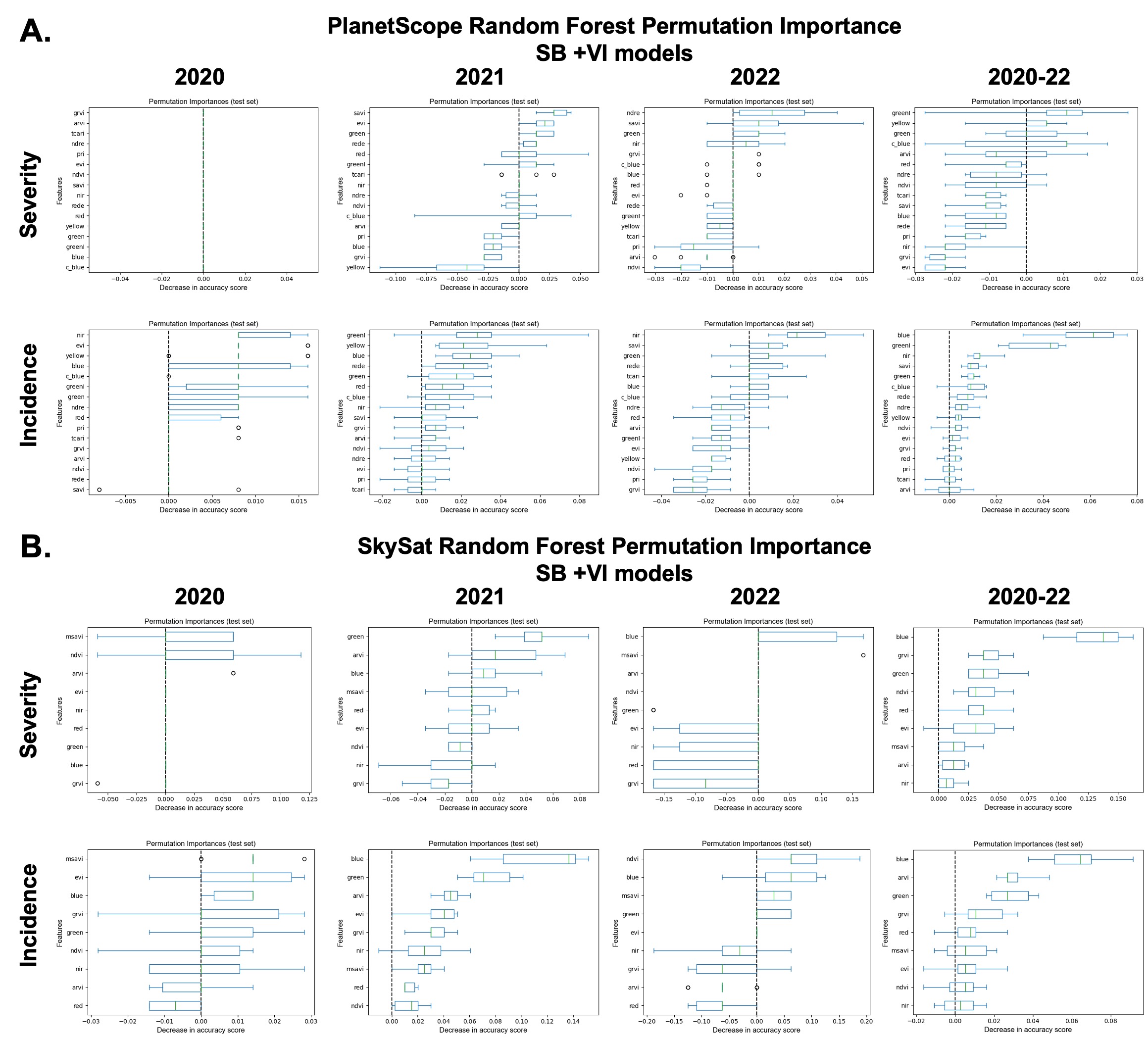
